# Clouds, oases for airborne microbes – Differential metagenomics/ metatranscriptomics analyses of cloudy and clear atmospheric situations

**DOI:** 10.1101/2023.12.14.571671

**Authors:** Raphaëlle Péguilhan, Florent Rossi, Muriel Joly, Engy Nasr, Bérénice Batut, François Enault, Barbara Ervens, Pierre Amato

## Abstract

Bacteria cells and fungal spores can aerosolize and remain suspended in the atmosphere for several days, exposed to water limitation, oxidation, and lack of nutrients. Using comparative metagenomics/metatranscriptomics, we show that clouds are associated with the activation of numerous metabolic functions in airborne microorganisms, including fungal spore germination. The whole phenomenon mirrors the rapid recovery of microbial activity in soils after rewetting by rain, known as the “Birch effect”. Insufficient nutrient resources in cloud droplets cause a famine that recycling cellular structures could alleviate. The recovery of metabolic activity by microorganisms in clouds could favor surface invasion upon deposition, but it may also compromise further survival upon cloud evaporation. In any case, clouds appear as floating biologically active aquatic systems.

**One-Sentence Summary:** Clouds activate metabolic processes in airborne microorganisms

## Microbes roam in the air

Microorganisms thriving on surfaces can detach and drift through the air for up to several days (*1*). There is a plethora of examples of air-dispersed microbes. Some attract attention when they are associated with sanitary threats to humans, crops or livestock, although these usually represent a small fraction of the total microbial biomass and diversity circulating in the natural atmosphere.

The low concentration and relatively short residence time of airborne particles make their characterization challenging, and microorganisms comprise only a small fraction of these. Microbiological studies are therefore usually limited to documenting microbial cell numbers and biodiversity, identifying probable sources, and at most listing the biological functions they carry and potentially spread. Airborne microbial assemblages consist of fungal spores, bacteria (isolated and aggregated), and fragments of biofilms in varying proportions and at concentrations generally ranging from ∼10^3^ to ∼10^6^ cell m^-3^ depending on meteorological, climatic and phenological conditions (*2, 3*). A wide range of environments from deserts to polar regions (*4, 5*) have been examined. As expected, microorganisms circulating in the air at proximity from the ground reflect the microbiota of the emitting surfaces and their temporal dynamics (*i*.*e*., seasonality of vegetation) (*6*–*8*). At high altitudes, the plumes from a variety of sources mix and result in diverse assemblages, despite very low biomass (∼10^2^-10^4^ cells.m^-3^) (*9*).

The role of the atmosphere as a carrier of (biological) material is therefore not in question, but how the functioning of living cells may be modulated during atmospheric transport remains largely unexplored. Once aerosolized, living microorganisms maintain viability and metabolic activity for some time (*10*–*13*), and they remain capable of responding to changing environmental conditions. For instance, bacteria (*Sphingomonas aerolata*) elevate ribosome numbers to activate their metabolic machinery in the presence of volatile organic nutrients (*14*). Meanwhile, survival and metabolic activity are challenged by low temperatures, limited water and nutrient availability and accessibility, and high levels of UV radiation and oxidants (*2*). The low water availability, in particular, is among the most limiting factors of biological processes in nature (*15*). Clouds, by offering liquid water to surviving airborne cells, could, thus, act as “oases” within the otherwise vast and hostile atmospheric environment.

## Clouds are floating aquatic systems

Clouds occupy ∼15% of the volume of the lower troposphere (*16*). These are air volumes where water condenses on the surface of aerosol particles, forming liquid droplets of a few micrometers in diameter and totalizing typically < ∼1 g/m(air)^3^. Chemical compounds from the gas and condensed phases dissolve into it, leading to an aqueous mixture of organic and inorganic compounds. Complex chemical processes occurring in the multiphase cloud system influence the composition of air masses (*16*–*19*), and *in-situ* microbiological processes may be affect organic compounds and oxidants (*20*–*22*). From the perspective of the microbiologist, cloud droplets can be considered as short-lived aquatic microhabitats providing microorganisms with liquid water and a range of dissolved nutrients at nano-to micro-molar concentration (carboxylic acids, amino acids, ammonium, nitrate, metals, etc.) (*23, 24*). Bulk cloud water thus offers conditions to thrive (*25, 26*). Moreover, clouds as the sources of precipitation act as an efficient way to the ground for airborne microorganisms (*27, 28*), where they may settle and establish.

## A new approach to addressing the guiding question

As far as the metabolic functioning of airborne microorganisms is concerned, most of our current knowledge is based on laboratory incubations of bulk samples (*20, 25*), or from isolated model strains injected into atmospheric simulation chambers (*13*), *i*.*e*. under conditions possibly not fully reflecting *in-situ* atmospheric conditions. Metagenomics and metatranscriptomics (*i*.*e*., analyses of whole DNA and RNA) provide snapshots of the biological processes taking place in a system. Over the last decade, the advent of high-throughput sequencing techniques stimulated such approaches and led to unprecedented insights into the functioning of microbiota in humans (*29, 30*), oceans (*31*), rivers (*32*), soil (*33*), and highly polluted environments (*34*). Clouds have been only very occasionally explored, revealing specific biological functioning compared to other biomes (*35*), but no information concerning specificities compared to the clear atmosphere. Using a novel untargeted approach, we examine here the possibility that clouds modulate and promote metabolic processes in airborne microorganisms.

Taking advantage of the atmospheric research station at the summit of Puy de Dôme Mountain (France, 1,465 m a.s.l.), often covered by clouds, we carried out a comparative analysis of the functioning of the natural aeromicrobiome in cloudy and clear atmospheres. This, to our knowledge, is unprecedented and required many experimental and analytical developments (*28, 35*–*37*). Various atmospheric conditions were assessed through a combined untargeted metagenomics/metatranscriptomics approach to reveal differences linked to the presence of clouds (see **Table 1** for information on sample acquisition and **Supplementary text** for details on the methods). Several high-flow rate impingers were deployed in parallel from a dedicated platform on the roof of the atmospheric station to collect airborne material directly into a nucleic acid preservation buffer, and capture information about the instantaneous functioning of living cells. Metagenomes (MGs) and metatranscriptomes (MTs) were obtained from shotgun sequencing of total nucleic acid extracts (**Table S1-S3**). Functional analyses were carried out from a composite gene catalog combining all the MGs, de-replicated, annotated, and used as a unique reference database for the whole study (see **Fig. S1-S2**) (*31*). Differential gene expression analysis (DEA) was then performed between clouds and clear atmosphere, and functions behind genes were categorized according to their corresponding Gene Ontology terms (GOs).

**Table 1.** Conditions of sample acquisition.

| SampleID | Sampling date<br>(dd/mm/yyyy) | Sampling<br>duration<br>(h) | Geographic<br>al origin of<br>the air<br>mass<sup>†</sup> | Boundary layer<br>height (min-max<br>[average]) (m<br>above sea level)<sup>‡</sup> | Position of<br>the sampling<br>site relative to<br>the boundary<br>layer | Temperat<br>ure (°C)<sup>§</sup> | Relative<br>humidity<br>(%)<sup>§</sup> | Wind speed<br>(m.s<sup>-1</sup>)<sup>§</sup> | Liquid water<br>content (LWC)<br>(g.m<sup>-3</sup>)<sup>§</sup> |
| --- | --- | --- | --- | --- | --- | --- | --- | --- | --- |
| **CLEAR**<br>**CONDITIONS** |  |  |  |  |  |  |  |  |  |
| 20200707AIR | 07/07/2020 | 6.5 | NW | 1268-1834 [1626] | In | 11.1 | 61 | 3.6 | < 0.01 |
| 20200708AIR | 08/07/2020 | 6.1 | NW | 623-1675 [1253] | In | 14.2 | 53 | 3.1 | < 0.01 |
| 20200709AIR | 09/07/2020 | 6.0 | N | 651-2377 [1487] | In | 20.3 | 48 | 3.4 | < 0.01 |
| 20200922AIR | 22/09/2020 | 5.9 | W | 665-1334 [972] | Out | 12.4 | 78 | 1.0 | < 0.01 |
| 20201118AIR | 18/11/2020 | 5.8 | W | 680-1142 [870] | Out | 14.1 | 41 | 6.4 | < 0.01 |
| 20201124AIR | 24/11/2020 | 6.0 | W | 644-740 [699] | Out | 8.6 | 50 | 3.4 | < 0.01 |
| **Minimum** | - | **5.8** | - | - | - | **8.6** | **41** | **1.0** | **< 0.01** |
| **Maximum** | - | **6.5** | - | - | - | **20.3** | **78** | **6.4** | **< 0.01** |
| **Median** | - | **6.0** | - | - | - | **13.3** | **52** | **3.4** | **< 0.01** |
| **Mean** | - | **6.1** | - | - | - | **13.5** | **55** | **3.5** | **< 0.01** |
| **Standard error** | - | **0.2** | - | - | - | **4.0** | **13** | **1.7** | - |
| **CLOUDS** |  |  |  |  |  |  |  |  |  |
| 20191002CLOUD | 02/10/2019 | 2.4 | NW | 1422-1505 [1465] | In | 6.5 | 100 | 3.0 | NA |
| 20191022CLOUD | 22/10/2019 | 6.4 | S | 698-957 [813] | Out | 5.7 | 100 | 8.7 | NA |
| 20200311CLOUD | 11/03/2020 | 4.1 | W | 964-1145 [1060] | Out | 5.0 | 100 | 7.4 | NA |
| 20200717CLOUD | 17/07/2020 | 3.3 | NW | 1271-1437 [1343] | Out | 10.1 | 100 | 1.6 | 0.08 |
| 20201016CLOUD | 16/10/2020 | 4.7 | NE | 917-1034 [958] | Out | 1.1 | 100 | 1.8 | 0.35 |
| 20201028CLOUD | 28/10/2020 | 6.0 | W | 1026-1529 [1269] | Out | 5.2 | 100 | 11.0 | 0.23 |
| 20201103CLOUD | 03/11/2020 | 3.5 | W | 1126-1593 [1390] | In | 2.2 | 100 | 8.7 | 0.06 |
| 20201110CLOUD | 10/11/2020 | 3.1 | SW | 691-1276 [1016] | Out | 5.9 | 100 | 2.5 | 0.07 |
| 20201119CLOUD | 19/11/2020 | 2.8 | W | 1207-1234 [1215] | Out | 0.3 | 100 | 7.7 | 0.11 |
| **Minimum** | - | **2.4** | - | - | - | **0.3** | **100** | **1.6** | **0.06** |
| **Maximum** | - | **6.4** | - | - | - | **10.1** | **100** | **11.0** | **0.35** |
| **Median** | - | **3.5** | - | - | - | **5.2** | **100** | **7.4** | **0.10** |
| **Mean** | - | **4.0** | - | - | - | **4.7** | **100** | **5.8** | **0.15** |
| **Standard error** | - | **1.4** | - | - | - | **3.0** | **0** | **3.6** | **0.11** |
| **P-value (Mann-Whitney test; clouds**<br>**vs clear conditions)** |  | **0.04\*** | - | - | - | **0.003\*\*** | **0.001\*\*** | **0.44** | **0.003\*\*** |

\*: significant p-value (< 0.05); \*\*: highly significant p-value (< 0.01); †: derived from 72-hour air mass backward trajectory, as detailed in Supplementary text; ‡: data extracted from ECMWF ERA5 model for the sampling period; §: average over the sampling period; NA: No data available.

## Not all microorganisms are equal before clouds

The datasets consist of sequences from Bacteria, Eukaryota, Archaea and viruses, with Eukaryota dominant in clouds and bacteria during clear weather (**Fig. S3**). Proteobacteria and Actinobacteria in Bacteria, and Ascomycota and Basidiomycota in Eukaryotes are prominent in all samples, as usual over continental vegetated areas (**Data S1**-**S2; Fig. S4-S5**) (*38, 39*). In both clouds and clear atmosphere, active taxa (*i*.*e*., exhibiting RNAs) represent ∼20% of the biodiversity present (*i*.*e*., detected in MGs) and tend to converge toward certain taxa (**Fig. S6**), which could result from selection processes linked with aerial transport.

Higher total relative RNA concentration in clouds as compared to cloud-free air points toward higher microbial metabolic activity (*31*) (**Fig. 1A; Table S1**). Yet, not all active taxa respond equally to the presence of condensed water, as shown by their representation in MTs with respect to MGs (abbreviated as RNA:DNA ratio, used as an indicator of the level of potential metabolic activity, (*40*)) (**Data S3; Fig. S7-S8**). Eukaryotes usually exhibit a higher RNA:DNA ratio than bacteria under both atmospheric conditions. Certain taxa of fungi have a higher ratio in clouds, such as *Botrytis* and *Colletotrichum*, suggesting higher metabolic activity, while others such as *Ustilago* -all common parasites of plants-have a higher ratio during clear conditions. In bacteria, this ratio is either unaffected by the presence of clouds, or elevated in some members of Gamma-Proteobacteria (*Pseudomonas*), Actinobacteria (*Rhodococcus*), and Alpha-Proteobacteria (*Sphingomonas*), in accordance with previous findings identifying active microorganisms in cloud water (*12*).

**Fig. 1.**
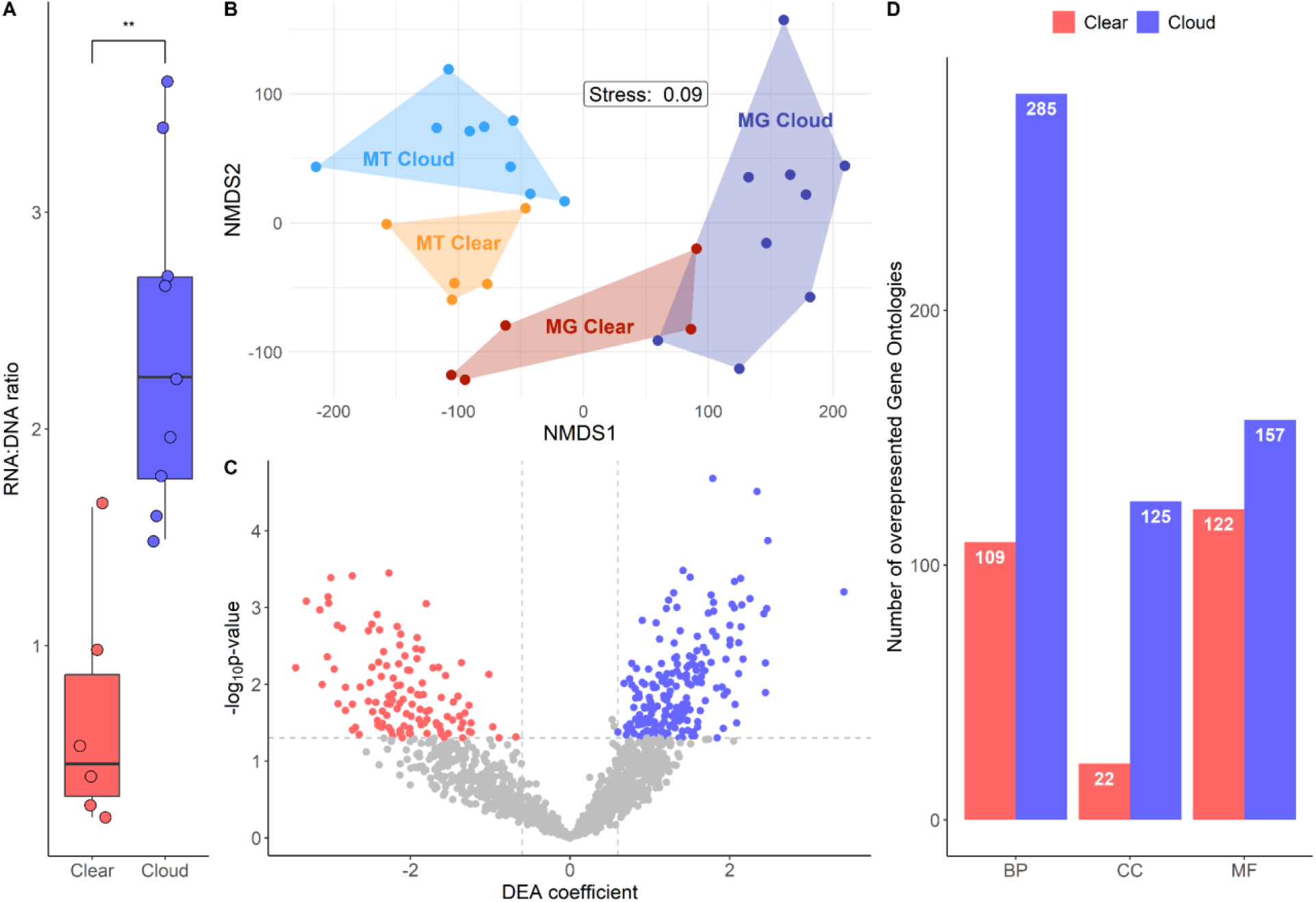
Comparative gene expression during cloudy and clear conditions: **(A)** RNA-to-DNA concentration ratio for clouds and clear atmospheric conditions; dots indicate individual samples, and boxplots display medians, 25^th^ and 75^th^ percentiles, and whiskers 1.5 interquartile ranges; ** indicates highly significant difference, highlighting the relatively higher total RNA concentration in clouds (Mann-Whitney test, p-value=0.004); **(B**) Non-Metric Multidimensional Scaling (NMDS) polygons based on the 21,046 functional gene entries in MGs and MTs, illustrating the distinction between MGs and MTs, and the differences between MTs during cloudy and clear conditions; **(C)** volcano plot representing differential gene expression between atmospheric conditions, with genes significantly overexpressed in and outside clouds in blue and red, respectively; grey dots represent genes whose expression level is not affected; dashed lines represent significance thresholds; DEA coefficient: Differential Expression Analysis (DEA) coefficient, from the MTXmodel R package (*78*); this illustrates that overexpression of distinct genes depends on the atmospheric condition; **(D)** numbers of overrepresented Gene Ontology terms (GOs) derived from gene annotations during cloudy and clear conditions, in blue and red, respectively, referring to their classification as Biological Processes (BP), Cellular Components (CC) and Molecular Functions (MF). A number of biological functions are overexpressed in clouds compared with clear conditions.

Typical atmospheric residence times of microbial cells are on the order of a few days (*1*), during which they may undergo ∼10 water evaporation-condensation cycles before precipitating (*41*). Whereas clouds may exist for several hours (*42*), individual cloud droplets evaporate after a few minutes (*43*). The lag-time after which microorganisms start recovering in soil upon rewetting shortens after repeated dry-wet cycles, due to selection processes toward the most responsive ones (*44*). In the air such rapid alternance of water availability could select as well for the most responsive microorganisms, like Proteobacteria and Actinobacteria (*45*), both indeed often dominating in airborne microbial assemblages and including numerous generalists with high metabolic flexibility (*46*).

## A “Birch effect” up in the sky

Comparative functional analysis between clouds and clear atmosphere reveals notable differences, and the data imply that clouds are associated with a resurgence of specific biological processes in airborne microbial cells (**Fig. 1B-D**). From a total of 21,046 genes detected in MGs, 488 (2.3%) can be qualified as “overexpressed”, *i*.*e*., significantly more represented in MTs than in MGs (**Data S4**), corresponding to a total of 1,005 distinct Gene Ontologies (GO) (**Data S5**). Eighty percent of those genes involve Eukaryotes, particularly Fungi. In Bacteria, nearly half of the genes overexpressed are associated with Gamma-Proteobacteria (**Fig. S9**). From these, a total of 320 genes (65.6% of the overexpressed genes) are significantly differentially expressed between clear and cloudy conditions (**Data S6-S7**). Eukaryotes contribute most overexpressed genes in clouds, and bacteria during clear conditions (**Fig. S10**). These relate to 820 GOs, on which our interpretations are based.

Several key metabolic functions are triggered in clouds, such as energy metabolism, transports of ions and carbohydrates, starvation, along with the down-regulation of oxidative stress and SOS responses (**Fig. 2-3, Fig. S11-S12)**. Differences are also notable regarding the expression of specific cellular responses to cloud conditions, such as response to osmotic stress, regulation of intracellular pH, and processes of autophagy and pexophagy (*i*.*e*., macropexophagy). This phenomenon mirrors the “Birch effect” that occurs in soils, whereby a burst of microbial activity is triggered in response to rewetting by rain (*47, 48*). Our data also reflect fungal spore germination, during which functions of cell protection give way to anabolic processes, within minutes (*49, 50*). In both cases, microbial growth and respiration are promoted by the sudden influx of water, associated with the release and solubilization of readily bioavailable organic compounds.

**Fig. 2.**
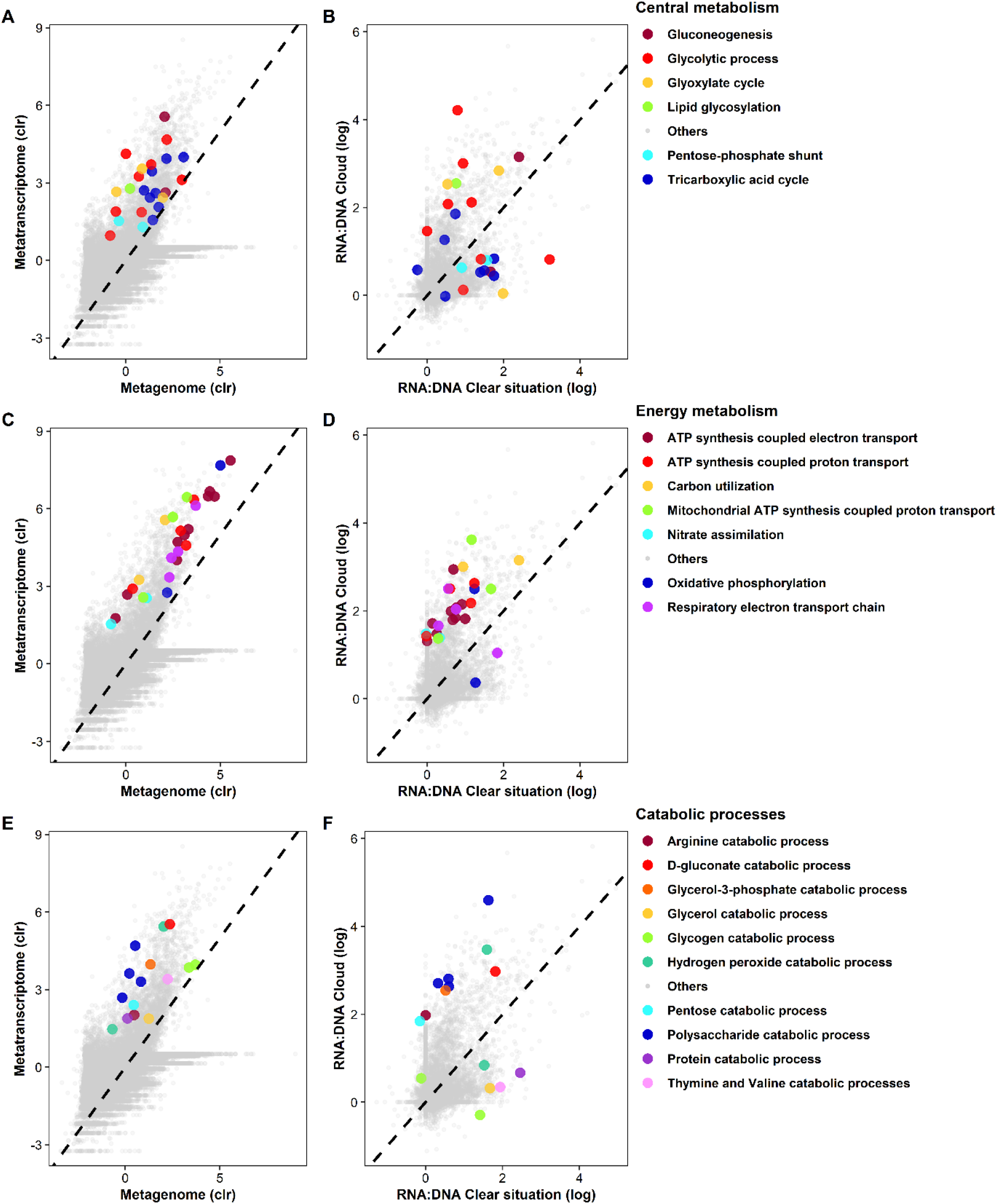
Representation of Gene Ontology terms (GOs) in MTs *versus* MGs, all samples considered (A; C; E), and functional expression levels, in clouds *versus* clear conditions based on their representation in MTs *versus* their corresponding MGs (termed as RNA:DNA) (B; D; F), for Biological Processes related to central metabolism (A, B), energy metabolism (C, D), and catabolic processes (E, F); clr: centered log-ratio transformation. Other GOs of interest are presented as Fig. S7 and S8.

**Fig. 3.**
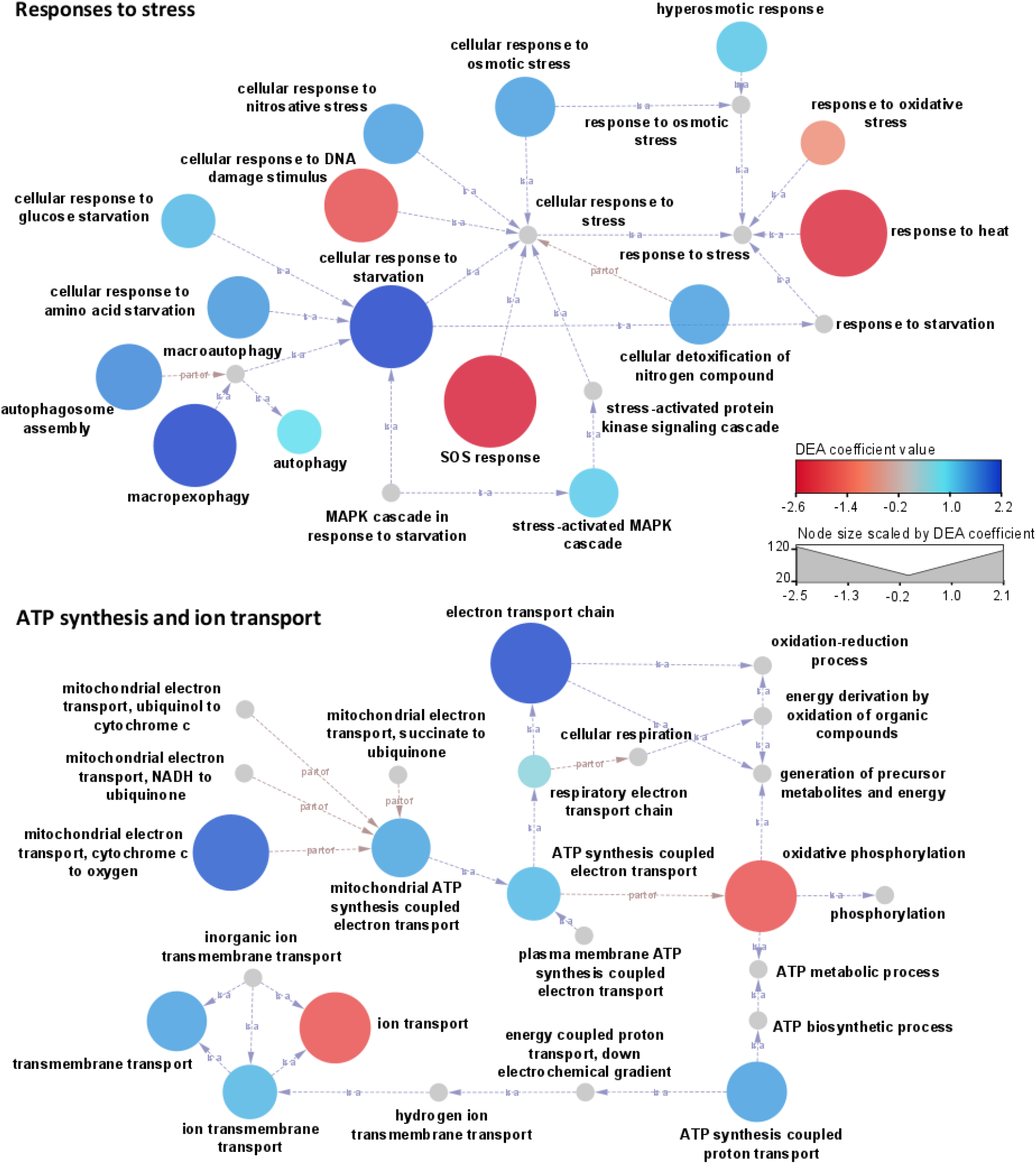
Gene Ontology terms (GOs) relationship networks linking different Biological Processes related to stress responses or ATP synthesis and ion transport, in clouds and clear atmosphere. The red-to-blue color scale represents the Differential Expression Analysis (DEA) coefficient, with negative values (red shades) indicating a significant overrepresentation in clear conditions, and positive values (blue shades) indicating overrepresentation in clouds. The size of the nodes is scaled by the absolute value of the corresponding DEA coefficient. Arrows indicate relationships between GOs (“*is a*” or “*part of*” as specified).

The “Birch Effect” in soils typically lasts for a few days, hence, much longer than the lifetime of individual cloud droplets (*43*). The metabolic modulations of airborne microorganisms depicted here should then be true for any cloud independently of its age. As in soils - but necessarily to a much smaller extent due to the low biomass - this phenomenon may lead to the release of gaseous biogenic compounds such as N_2_O through aerobic ammonium oxidation, *i*.*e*. nitrification (*51*), along with CO_2_ from respiration.

Fungal spores are dispersal propagules designed for (aerial) spreading over long distances (*52*) until they reach favorable conditions to germinate. While dormant, functions of cell protection against osmotic stress, heat, and oxidants are expressed. In the presence of water, the germination process starts within minutes, involving mitogen-activated protein kinases (MAPK) cascade signaling, and leads to a switch toward the activation of central metabolic functions involved in energy production and biosynthesis (*50*). Our observations strongly concur with such a sequence, and it is reasonable to assert that airborne fungal spores do germinate in clouds.

Many functions related to starvation and autophagy suggest that the nutritive requirements in clouds are not fully satisfied. Bulk cloud water typically contains enough diverse nutrients to sustain metabolic processes and some extent of microbial growth (*22, 25*). However, statistically only ∼1 out of 10,000 droplets contains a microbial cell that potentially causes a very efficient and rapid depletion of nutrients in these small volumes (∼10^−6^ μL for 20 μm diameter droplets) (*21*). Autophagy processes can help compensating for the needs, and contribute initiating the synthesis of the appressorium in fungi, a structure designed to invade host cells in parasites and symbionts (*53*). Peroxisomes, organelles involved in the detoxification of oxidants in eukaryotes, are targeted in particular by autophagy processes (pexophagy), which could compromise the chances of survival toward further aerial transport.

Water limitation is a great challenge that many microorganisms face in their common natural habitat. This is particularly true on surfaces interfacing with air, where humidity conditions can vary widely within short periods of time. Cloud-free air was collected at an average relative humidity (rH) ranging from 41% to 78%, levels of water activity barely compatible with biological processes (*54*). We found no correlation of any biological process with rH outside clouds. In both cloudy and clear conditions, central metabolic and catabolic functions such as glycolysis, tricarboxylic acid cycle, and pentose phosphate shunt are detected, as well as biosynthetic processes directed toward nitrogen-containing compounds, typically amino-acids, indicative of on-going translational activity, *i*.*e*. protein synthesis (**Fig. S11**). Functions of glutamine and glycine synthesis in particular are preferred in clouds, whereas that of lysine occurs in any condition, from aminoadipic acid in clouds or from diaminopimelic acid in clear air. These latter pathways participate in peptidoglycan synthesis in bacteria (*55, 56*), and might be responses to osmotic variations and attempts to multiply. Although not evaluable here, it is conceivable that some biomass production occurs in clouds, but it is necessarily constrained by short residence time and low temperatures (*57*). Yet, given that microbial cells are likely to encounter multiple cloud cycles during their atmospheric journey, the diversity of airborne microbial assemblages could be significantly modified between emission and deposition.

Whether or not recovery of metabolic activity, germination or even multiplication in clouds provides any advantage to airborne microorganisms remains unclear. Multiplying appears by essence advantageous and may favor the most responsive microorganisms to invade surfaces or hosts upon deposition (*58*). In turn, triggering germination and sacrificing essential cellular structures while conditions may soon become inhospitable could compromise future chances of survival and further dispersion in the likely event where the cloud evaporates, instead of precipitating.

References 59 to 83 are only included in SM.

## Acknowledgments

We tank OPGC’s facility for running the PUY atmospheric station and sharing meteorological data, and for logistical help during sampling. We are also grateful to the computational center of Clermont Auvergne University (Mesocentre) and the network Auvergne Bioinformatique for providing computational power and excellent support with software deployment in the Galaxy environment.

## Author contributions

Conceptualization: PA

Methodology: RP, FR, FE, BB, EN

Investigation: RP, PA Visualization: RP

Funding acquisition: PA, BE

Supervision: PA

Writing – original draft: RP, PA

Writing – review & editing: PA, BE

## Data and materials availability

Raw sequencing MGs and MTs data are available as fastq.gz files through the European Nucleotide Archive at EBI, under the project accession PRJEB54740, samples ERR9966616 to ERR9966643.

## Supplementary Materials

Materials and methods Figs. S1 to S12

Tables S1 to S3

References (*59-83*) Captions for Data S1 to S7

Data S1 to S7 (as separate electronic files)

