## Supplementary Materials for "Clouds, oases for airborne microbes – Differential metagenomics/ metatranscriptomics analyses of cloudy and clear atmospheric situations"

† Now at : IPREM – Institut des Sciences Analytiques et de Physico-Chimie pour l'Environnement et les Matériaux (IPREM) ; Pau, France.

‡ Now at : Département de Biochimie, de Microbiologie et de Bio-informatique, Faculté des Sciences et de Génie, Université Laval ; Québec, Canada.

**This supplement includes:**

Supplementary Text (Materials and methods)

Figures S1 to S12

Tables S1 to S3

Captions for Data S1 to S7

Data S1 to S7 (as separate electronic files)

Supplementary text:

Materials and Methods

Sample collection

Samples were collected from the summit of Puy de Dôme Mountain (PUY; 1,465 m a.s.l., 45.772° N, 2.9655° E, France) located ~400 km East of the Atlantic Ocean and ~300 km North of the Mediterranean Sea. This rural area is surrounded mainly by deciduous forests and pastoral landscapes, with the urban agglomeration of Clermont-Ferrand located ~8 km East. The PUY station is mainly exposed to winds from North and West (*24*, *59*). This station is part of the Cézeaux-Aulnat-Opme-Puy-de-Dôme (CO-PDD) instrumented platform for atmospheric research (*60*) and is fully equipped for on-site sample processing and conditioning.

A total of nine cloud and six clear atmosphere samples were collected in 2019 and 2020 for periods of ~two to ~six consecutive hours (**Table S1**). In both situations, several high-flow-rate impingers (HFRi; model DS6, Kärcher SAS, Bonneuil-sur-Marne, France) sampling with an air-flow rate of 2 m^3^/min (*37*) were deployed in parallel and treated as individual replicates. Nucleic acid analyses were carried out from dedicated HFRis, filled with 1.7 L of 0.5 X Nucleic Acid Preservation (NAP) buffer solution (*61*, *62*) as the collection liquid (see below) in the case of clear atmosphere, or 850 mL of 1 X NAP in the case of cloud. Complementary biological and chemical analyses requiring no saline solution were carried out from samples collected in parallel, using another HFRi filled with 1.7 L of autoclaved ultrapure water for clear atmosphere using, or a cloud droplet impactor sterilized by autoclave in the case of cloud as in (*63*). The loss of collection liquid in HFRIs due to evaporation during clear atmosphere sampling was compensated with autoclaved ultrapure water every hour based on HFRI sampling tank’s weight. Sampling blanks were performed at each sampling occasion. The control for nucleic acid-based analyses consisted of a volume of NAP buffer left in a HFRI sampling tank for >10 min, with the sampler set to “off” and the inlet closed. The controls for other analyses consisted of samples of the water used as the sampling liquid, taken from the collection tank of the sampler just before sampling. Samples and blanks were processed using the PUY station’s microbiology facility, within a laminar flow hood previously exposed to UV light for 15 min.

Immediately after sampling, the NAP buffer from each HFRi was individually filtered through 0.22 µm porosity mixed cellulose esters (MCE) filters (47 mm diameter; ClearLine, ref. 0421A00023) using individual sterile Nalgene filtration units. Filters were rolled using sterile forceps and placed into 5 mL Type A Bead-tubes (Macherey-Nagel, ref. 740799.50). A volume of 1,200 µL MWA1 lysis buffer (Macherey-Nagel ref. 740799.50) was then added to each tube, and a bead-beating step of 10 minutes was performed using a Genie 2 vortex set at maximum speed and the vortex adapter recommended in the nucleic acid extraction kit used (Macherey-Nagel NucleoMag^®^ DNA/RNA Water kit for water and air sample). Filters and lysates were finally stored at -80°C in the bead tubes until further processing.

Meteorological data and backward trajectory plots

Meteorological variables were monitored at the PUY meteorological station: temperature, relative humidity, liquid water content, wind speed and direction (<https://www.opgc.fr/data-center/public/data/copdd/pdd>). The boundary layer height (BLH) was extracted from ECMWF ERA5 global reanalysis (<https://www.ecmwf.int/en/forecasts/datasets/reanalysis-datasets/era5>) (*64*). The geographical origin of the air masses was derived from 72-hour backward trajectories computed using the CAT trajectory model (*60*), which uses dynamical fields extracted from the ERA-5 meteorological data archive with a spatial resolution of 0.125° for the present work. This tool allowed to estimate percentages of air mass trajectory points in each of the 8 direction sectors (*59*).

Nucleic acid preservation (NAP) buffer

NAP buffer was prepared according to the protocol described in (*61*). This buffer protects nucleic acids, particularly RNA, from degradation, while fixing the samples directly at the time of sampling. The buffer is composed of 0.019 M of ethylenediaminetetra-acetic acid (EDTA) from disodium salt dihydrate, 0.018 M of citrate from trisodium salt dihydrate, 3.8 M of ammonium sulfate, and sulfuric acid to adjust pH at 5.2. NAP buffer was filtered through GF/F and then autoclaved before use. Large volumes of 1X NAP buffer were prepared to match the needs imposed by the sampling strategy, by series of 10 L each; 0.5X NAP buffer was prepared by adding 1 volume of deionized water before autoclaving.

Nucleic acid extraction and shotgun sequencing

Total DNA and RNA were extracted simultaneously from individual MCE filters using NucleoMag® DNA/RNA Water kit (Macherey-Nagel, Hoerdt, France). All facilities were previously treated with RNase away spray solution (Thermo Scientific; Waltham, USA). For DNA extraction, 600 µL of lysate was processed following the extraction kit protocol adapted for 47 mm filter membranes, including RNA removal by the addition of 1:50 volume of RNase A (12 mg/mL, stock solution from Macherey-Nagel). DNA was eluted into 50 µL of DNase-free H_2_O after 5 min of incubation at 56°C, then quantified by fluorescence using the Quant-iT™ PicoGreen® dsDNA kit (Invitrogen; Thermo Fisher Scientific, Waltham, MA USA). For RNA extraction, the remaining 600 µL of lysate were processed following the protocols for 47 mm filter membrane, including DNA removal by the addition of 1:7 volumes of reconstructed rDNase (*cf* kit standard protocol). Total RNAs were eluted into 30 µL of RNase-free H_2_O after 10 min of incubation at room temperature.

Individual DNA or RNA extracts from the same sampling events were pooled and 30 µL of each were transferred to a subcontractor (GenoScreen, Lille, France) for further processing of RNAs (quantification, reverse-transcription to cDNAs), and shotgun sequencing of the metagenomes (from DNAs) and metatranscriptomes (from cDNAs) by Illumina HiSeq 2*150 bp. A first sample (20191022CLOUD) was used to check the feasibility of the approach, adjust the sequencing depth, and elaborate bioinformatics workflows; it was thus deeply sequenced (~200 M reads). The other samples were sequenced with a lower sequencing depth (40 - 60 M reads per sample). Due to the low concentrations of DNA or cDNAs in the sequencing libraries, several samples were pre-concentrated from additional volumes of the corresponding extracts (20 µL re-eluted into 8 µL).

Elaboration of the bioinformatics workflow for differential metagenomics/metatranscriptomics analyses

The bioinformatic strategy includes a dual differential analysis: MT against their corresponding MG, to investigate gene expression levels in each sample considering their abundance, and cloudy against non-cloudy conditions, to detect the specificities of gene expression. Thus, the objective was to obtain statistics of [MT_reads_/MG_reads_]_clear_ *versus* [MT_reads_/MG_reads_]_cloud_, with MT_reads_ and MG_reads_ the read counts in MT and MG, respectively, at the sequence, gene, taxon or Gene Ontology term (GO) level.

The bioinformatics workflow (BiW; **Figure S1**) was elaborated by assembling relevant existing tools on a Galaxy instance (*65*) deployed by the bioinformatics facility hosted by the University (Auvergne BioInformatique, AuBI), and the regional calculation cluster Mesocentre Clermont Auvergne. This includes large databases, *i.e.* the most adapted to environmental samples, and it enables both the processing of MGs and their corresponding MTs to standardize read counts data in the absence of absolute reference metagenome, and statistical analysis for comparing environmental situations. Briefly, the BiW contains the typical following steps: preprocessing of the sequences (quality control, trimming, assembly, etc.), taxonomic affiliation, construction of a catalog including all sequences, cleaned for redundancy, gene prediction and their functional and taxonomic annotations, mapping of each sample dataset toward the annotated catalog, statistical differential expression analysis (DEA) of MTs against their corresponding MGs to detect overexpressed genes and the associated taxa and function in every sample, and DEA of functional gene expression in cloudy *versus* clear conditions, in order to reveal differences between atmospheric conditions.

Raw MGs contained approximately between 30 M and 260 M reads, and raw MTs from 65 M to 195 M reads (**Tables S2-S3**), both with an average read size of 150 bp. The preprocessing step of our BiW consisted of sequences of FastQC (v 0.72) (*66*) for quality control (QC) analysis, Trimmomatic (v 0.36.6) (*67*), to filter and trim erroneous reads, with an initial ILLUMINACLIP step for remaining Nextera (paired-reads) adapters, and with a sliding window of 10:30, a reads min length of 100 bp and the leading and trailing parameters with a quality threshold at 30; this step removed between 26% and 37 % of the reads in MGs and between 20 to 49 % of the reads in MTs. Then SortMeRNA (v 2.1b.6) (*68*) to filter and recover the rRNA gene reads in separated files, with the default parameters, “paired-out” option and all the available databases. The rejected files (non-rRNA gene reads) were kept for downstream functional analyses. Proportions of rRNA gene reads in MG and MT datasets were typically between 1 and 2 % and between 80 and 94%, respectively, except for one sample (20201124AIR), whose MT inconsistently contained 12 % of reads annotated as rRNA genes, and this sample was thus removed from the analysis. One MT sample (20201103CLOUD) contained only 40 % of rRNA gene reads, but further analysis revealed that this was caused by a high amount of human sequences, which represented 90% of the non-rRNA reads. The presence of Human DNA in the atmosphere could be a topic of interest, but this is beyond the scope of our study. Human reads were filtered from the rejected files from SortMeRNA (non-rRNA gene reads) using Bowtie2 (v 2.4.2) (*69*), against the NCBI *Homo sapiens* genome “hg38_2021-5-18” with default parameters. Human sequences represented 0.01-0.71 % of the total trim MG sequences and 0.2-1.95 % of the total trimmed MT sequences (except in one sample, 90% as mentioned).

Taxonomic affiliations were obtained from whole MG and MT datasets using Kraken2 (v 2.1.1) (*70*) against the “PlusPF” Kraken database (as of 2021-1-27) including known archaeal, bacterial, viral plasmid, human, protozoan, and fungal genomes. This step was performed on trimmed files before rRNA gene sorting (SortMeRNA) and human sequence filtering (Bowtie2) steps. Kraken2 was used with a confidence score threshold of 0.1 and with the “report” and “report-zero-counts” options. In total, 1.2-4 % of the MG reads and 63-90% of the MT reads were affiliated, with 2.3-28.8 % and 0.02-0.7 % of the reads, respectively, affiliated to Human. Human reads were removed from the taxonomy datasets.

Since no reference metagenome or gene database is available for atmospheric environments, a catalog of non-redundant genes was constructed here by aggregating genes from all MGs, and then used as a reference database for the analysis of all samples. To this end, each MG was first *de novo* assembled using MEGAHIT (v 1.1.3.5) (*71*), considering only non-rRNA gene reads. Default parameters were used with a minimum length for contigs of 500 bp. Depending on the sample, the number of assembled contigs ranged from ~43,000 to ~495,000, representing a total of 2,832,534 contigs. The maximum size of the contigs was between 21,000 and 200,000 bp, with a mean size between ~750 and 1,010 bp depending on the sample. Genes were then predicted from all these contigs using MetaGeneAnnotator (v 1.0.0) (*72*). The “MetaGenomic” option was used, with BED format as the output file. From the total 2,832,534 contigs, a total of 3,168,750 genes were predicted. Finally, in order to prevent redundancy, gene sequences were clustered at 95% identity using CD-HIT (v 4.8.1) (*73*, *74*) and only the representative sequence of each cluster was kept. Only sequences >100 bp, and exhibiting >90 % identity with a reference sequence were kept. A total of 1,067,351 non-redundant genes were finally retained, with a length ranging from 100 to 22,065 bp and an average of 330 bp.

Functional annotation was performed on the genes catalog using DIAMOND (v 2.0.8.0) (*75*) in the “blastx” mode and default parameters and the UniProtKB Swiss-Prot functional gene database (as of 2021_03) (*76*). A total of 163,057 genes were annotated representing 40,264 unique UniProtKB entries.

Approximately 15% of the genes in the catalog were annotated, representing ~5% of the total predicted genes, which indicates high proportions of undocumented genes in the samples. A total of about 7% of the total entries in the UniProtKB Swiss-Prot database (566,996 entries in March 2022) were represented in the datasets, which illustrates the high biological diversity circulating in the atmosphere. A few UniprotKB entries were overrepresented in the gene catalog, mainly retroviruses and transposons from eukaryotes. The vast majority of the annotated genes (91.5 %) were associated with Eukaryotes, ~60 % with fungi, 23.6 % with viridiplantae and 14.3 % with metazoa. The remaining consisted of bacterial (7.6 %), viral (0.6 %) and archaeal genes (0.3 %) (**Figure S2**).

Non-rRNA gene sequences from all MGs and MTs were finally mapped to the gene catalog to obtain read counts in samples, using BWA-MEM (v 0.7.17.1) (*77*). Default parameters were used. The percentage of properly mapped reads mapped against the gene catalog was between ~4 and ~17 % for MGs and between ~3 and ~10 % for MTs (**Tables S2-S3**). Only genes with >10 mapped sequences in MGs were retained, and the count tables for MGs and MTs were filtered to remove gene IDs affiliated to “Embryophytes” and “Metazoa” and focus on microbial genes. Finally, 21,046 genes (over 40,264) remained for downstream analyses.

Data normalization and differential functional expression analysis

The normalization and differential expression analysis (DEA) were performed using the R package MTXmodel (R v4.0.3; MTXmodel v1.5.1) (*78*). This was run using the options: no transformation, clr (centered-log ratio) normalization, LM analysis method, BH correction method, “EnvType” as fixed effect (*i.e.*, cloudy or clear atmosphere), min abundance at 0.0001, min prevalence at 0.5, max significance at 0.25 and input of DNA data (MG count table). DEA gives coefficients of relative expression based on the fixed effect. Here, the environment type “cloud” was used as the reference, therefore positive coefficients (coeff) indicate that the feature (taxonomic group or gene) is significantly more represented in clouds, while negative coefficient indicates overrepresentation in clear conditions, with higher absolute values for higher representations. DEA was also performed independently from atmospheric conditions to differentiate MTs from MGs. The input files consisted of MT and MG datasets, the fixed effect was “MT or MG” and there was no “DNA data” input for standardization. Here, positive DEA coefficients identified the features significantly overrepresented in MTs.

Annotated sequences were grouped according to their corresponding Gene Ontology terms (GOs) (*79*, *80*), for Cellular Component, Molecular Function and Biological Process categories. REVIGO (*81*) was used to summarize and visualize GO lists.

Finally, the log ratios of the number of reads associated with a gene, taxon or GO in a MT dataset to that in the corresponding MG (abbreviated as RNA:DNA log ratios) were calculated using data normalized to total counts. RNA:DNA ratios are commonly used as an appraisal of the relative level of metabolic activity, with higher ratios indicating potentially higher metabolic activity (*40*, *82*).

Data visualization

Data visualizations were designed with the following R packages: *ggplot2* (v3.4.1), *ggrepel* (v0.9.3), *ggsignif* (v0.6.4), *ggdendro* (v0.1.23), *factoextra* (v1.0.7), *gridExtra* (v2.3), *vegan* (v2.6-4), *pheatmap* (v1.0.12). Relationship networks for GOs were generated using OLSVis (*83*) and Cytoscape (v3.9.1).

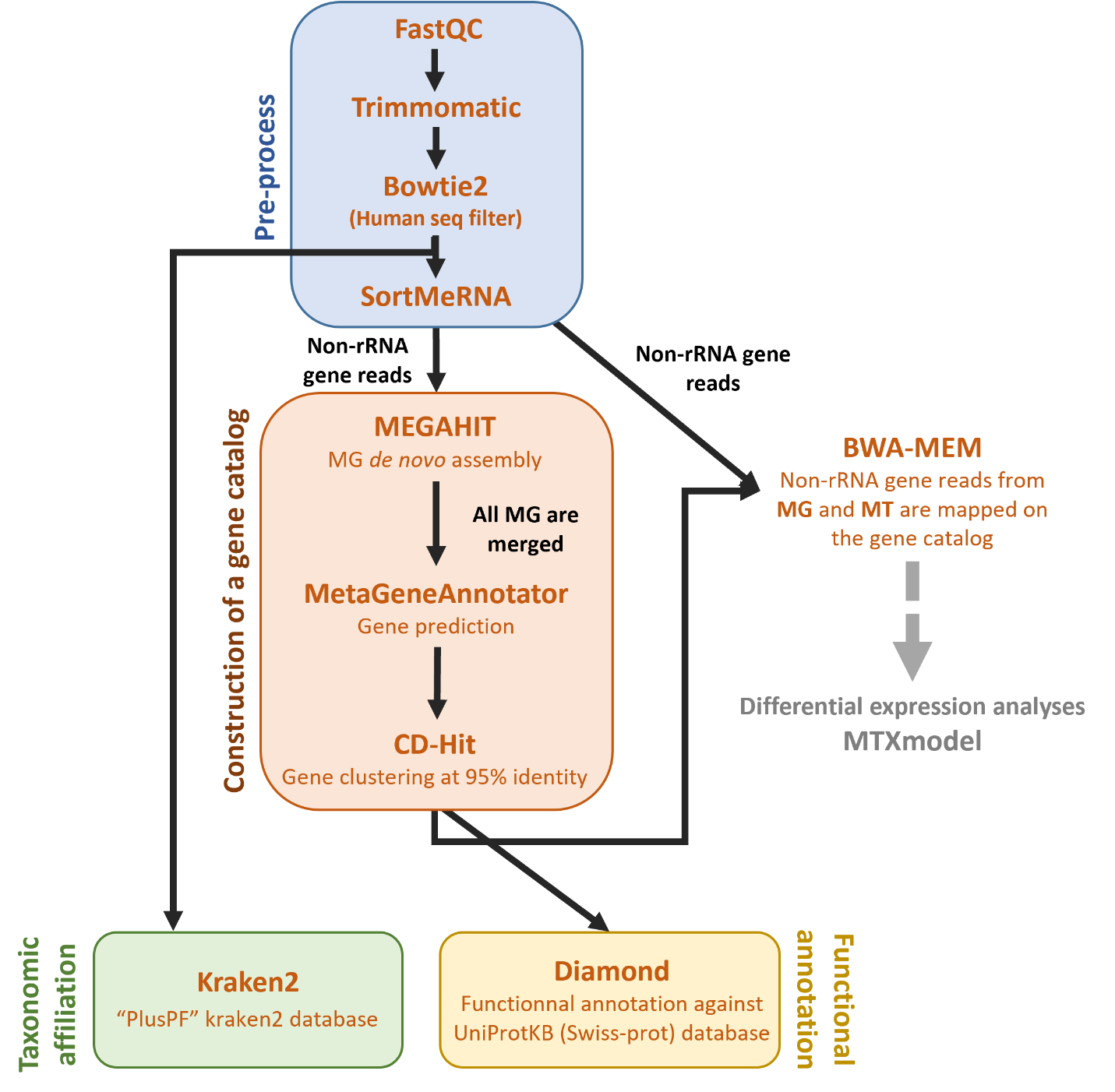

Fig. S1.

Main steps of the bioinformatics workflow elaborated for the analyses of combined MGs and MTs. This uses the following tools, run on Galaxy: FastQC (v0.72), Trimmomatic (v0.36.6), Bowtie2 (v2.4.2), SortMeRNA (v2.1b.6), MEGAHIT (v1.1.3.5), MetaGeneAnnotator (v1.0.0), CD-Hit (v4.8.1), Kraken2 (v2.11), Diamond (v2.0.8.0), BWA-MEM (v0.7.17.1), and the MTXmodel R package (v1.5.1) (see supplementary text for the corresponding references).

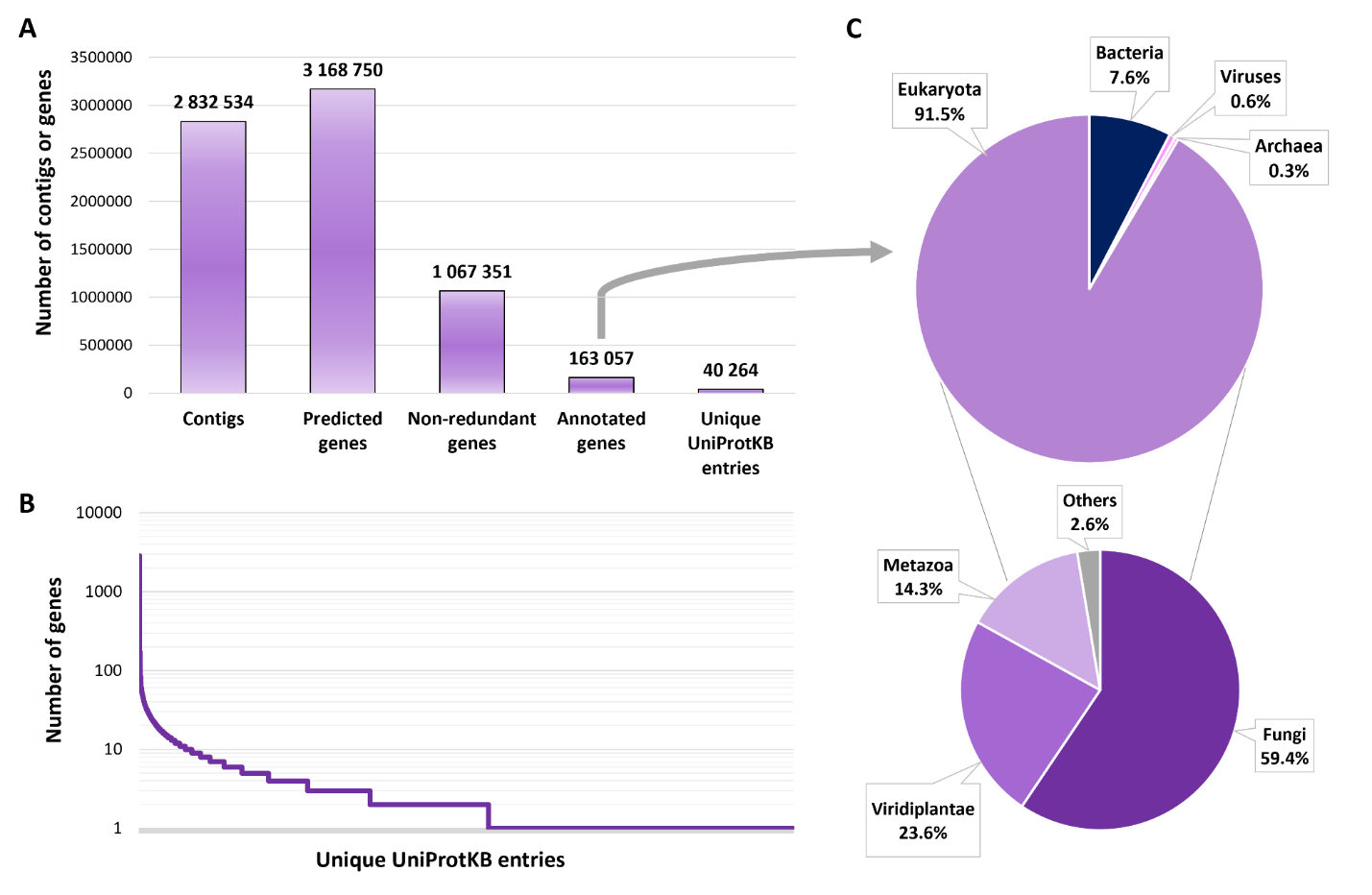

Fig. S2.

Content of the catalog of genes used as a reference, constructed by merging all MGs. A: Contigs or genes numbers at each step of the gene catalog construction, from left to right; B: Rank-abundance curve representing the number of genes associated with unique UniProtKB entries; C: Taxonomic affiliations associated with annotated genes; Eukaryota taxa are specified in the lower pie-chart.

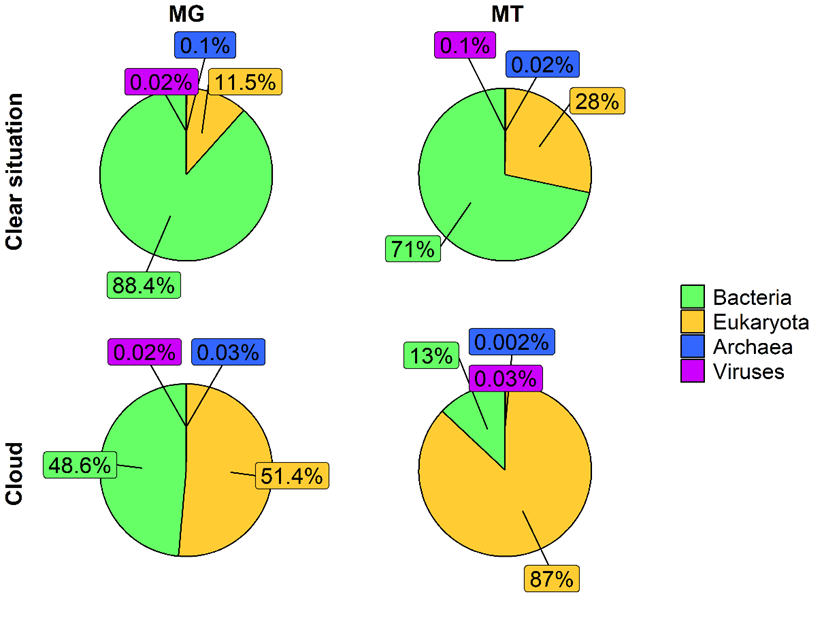

Fig. S3.

Average proportions of reads associated with Bacteria, Eukaryota, Archaea and Viruses in MGs and MTs during cloudy and clear conditions.

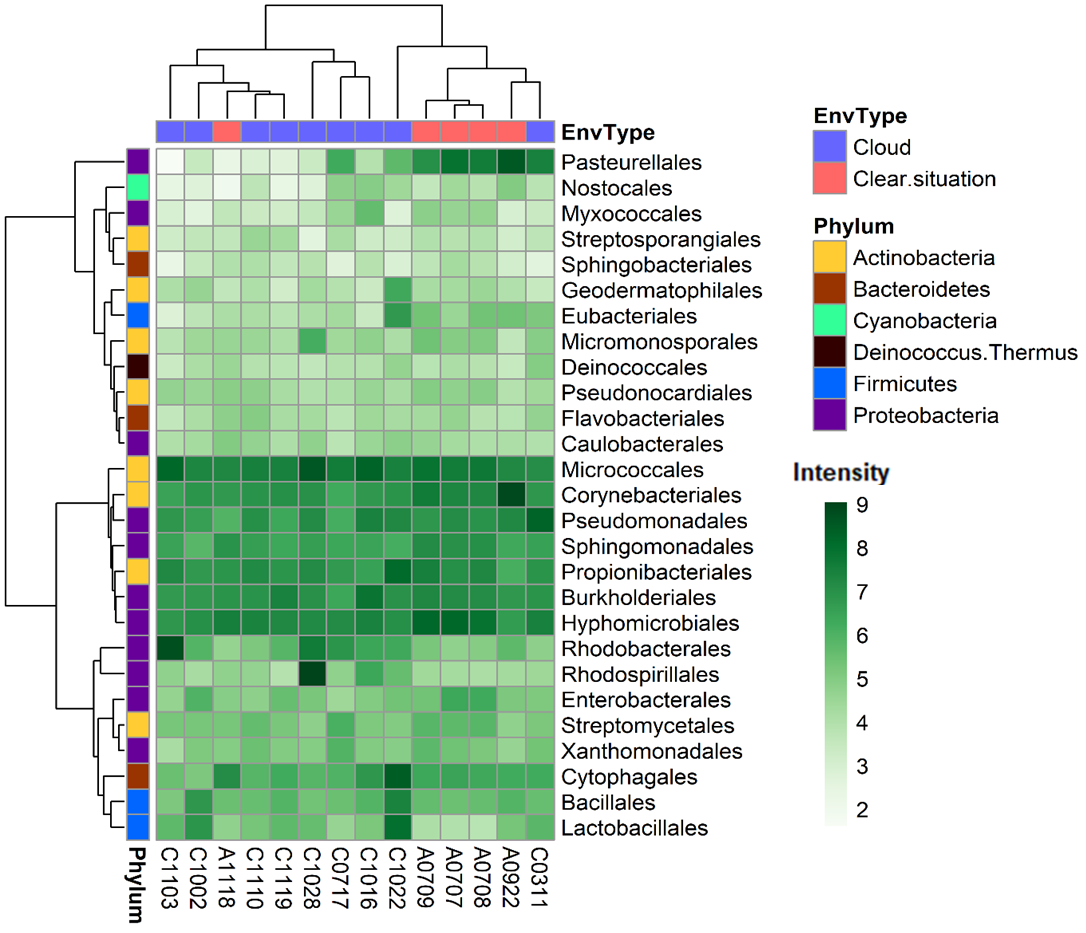

Fig. S4.

Distribution of the most abundant bacterial orders in MGs, and corresponding hierarchical clusterings (Ward’s method, “ward.D2”). The intensity scale depicts centered-log ratio (clr) abundances. EnvType: environment type. The samples are named as follows: “A” for clear conditions (air) or “C” for cloud, followed by the sampling date in the format “mmdd” (month and day).

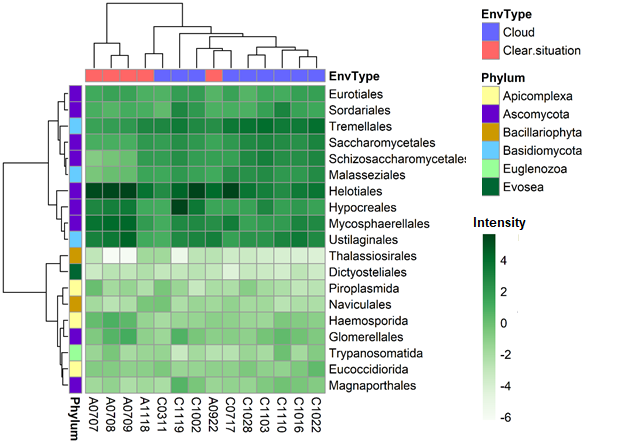

Fig. S5.

Distribution of eukaryotic orders in MGs, and corresponding hierarchical clusterings (Ward’s method, “ward.D2”). The intensity scale depicts centered-log ratio (clr) abundances. EnvType: environment type. The samples are named as follows: “A” for clear conditions (air) or “C” for cloud, followed by the sampling date in the format “mmdd” (month and day).

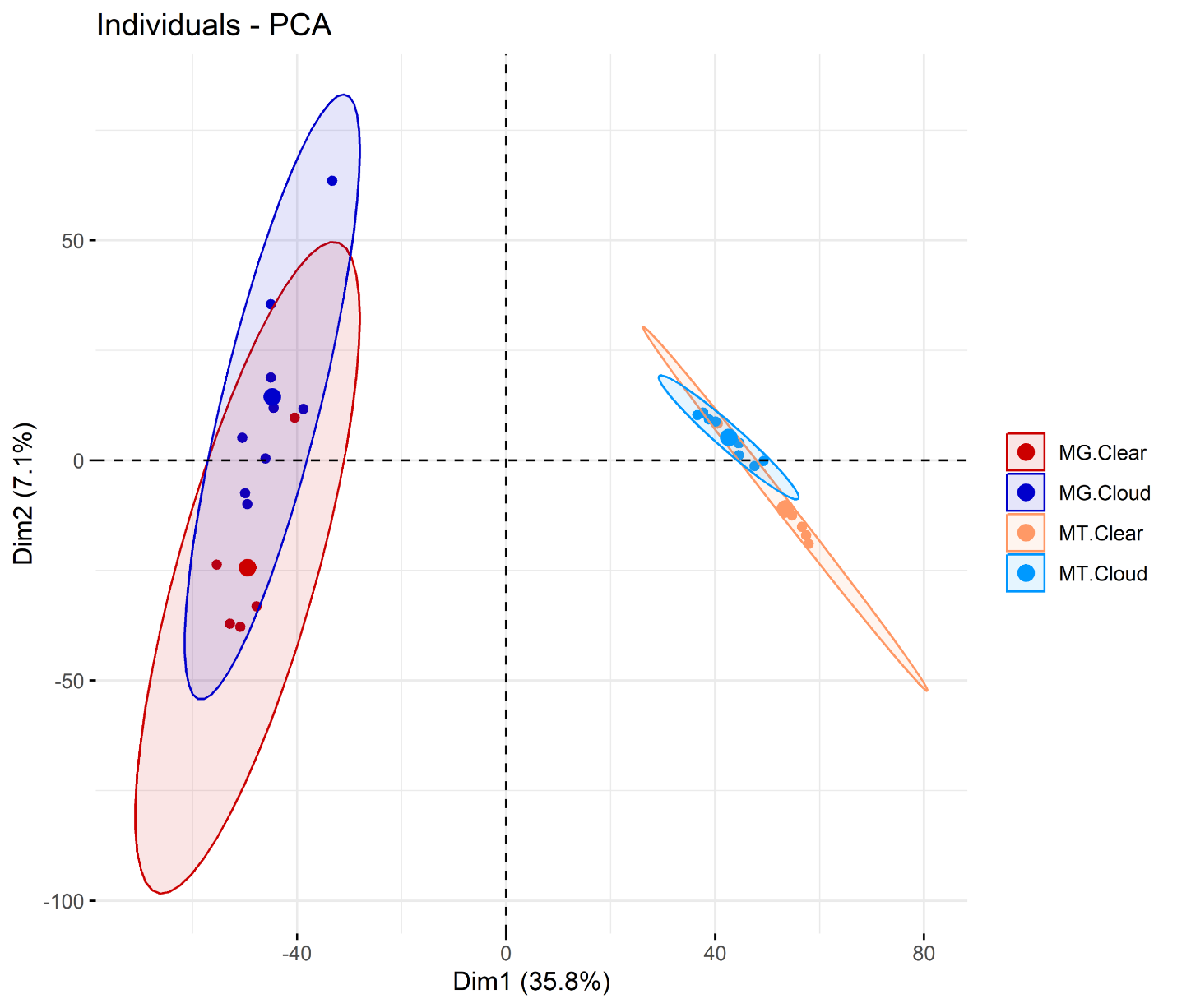
Fig. S6.

Principal component analysis (PCA) representing taxonomy distribution in MGs and MTs for cloudy and clear atmospheric conditions, based on 6,373 taxa. Count data were centered-log ratio (clr) transformed.

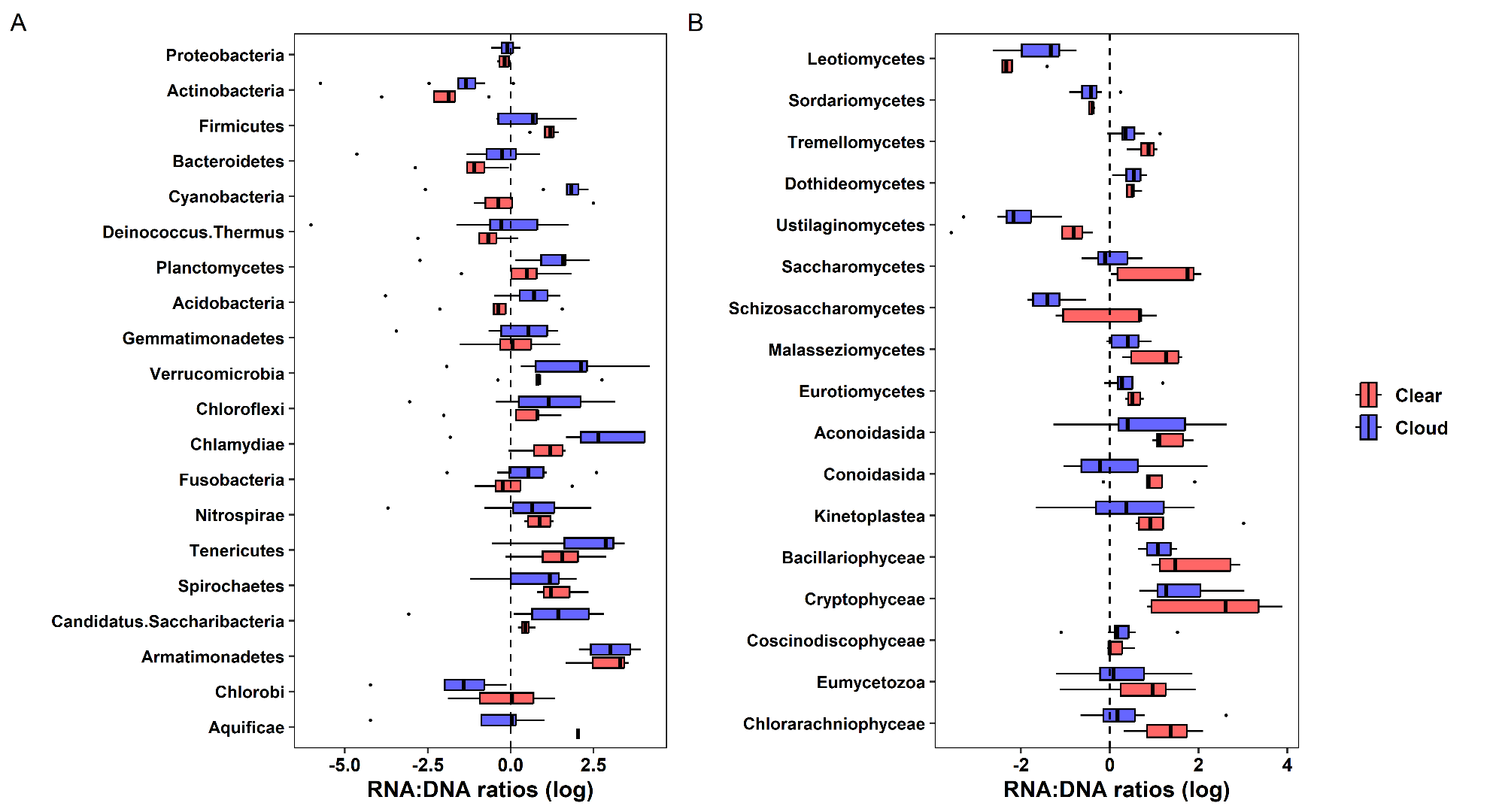

Fig. S7.

Relative representation of reads affiliated with (A) the 20 most represented phyla of bacteria and (B) classes of eukaryotes in MGs *versus* MTs for cloudy and clear atmospheric conditions. Taxa are ordered from top to bottom in descending order of the number of reads in MGs.

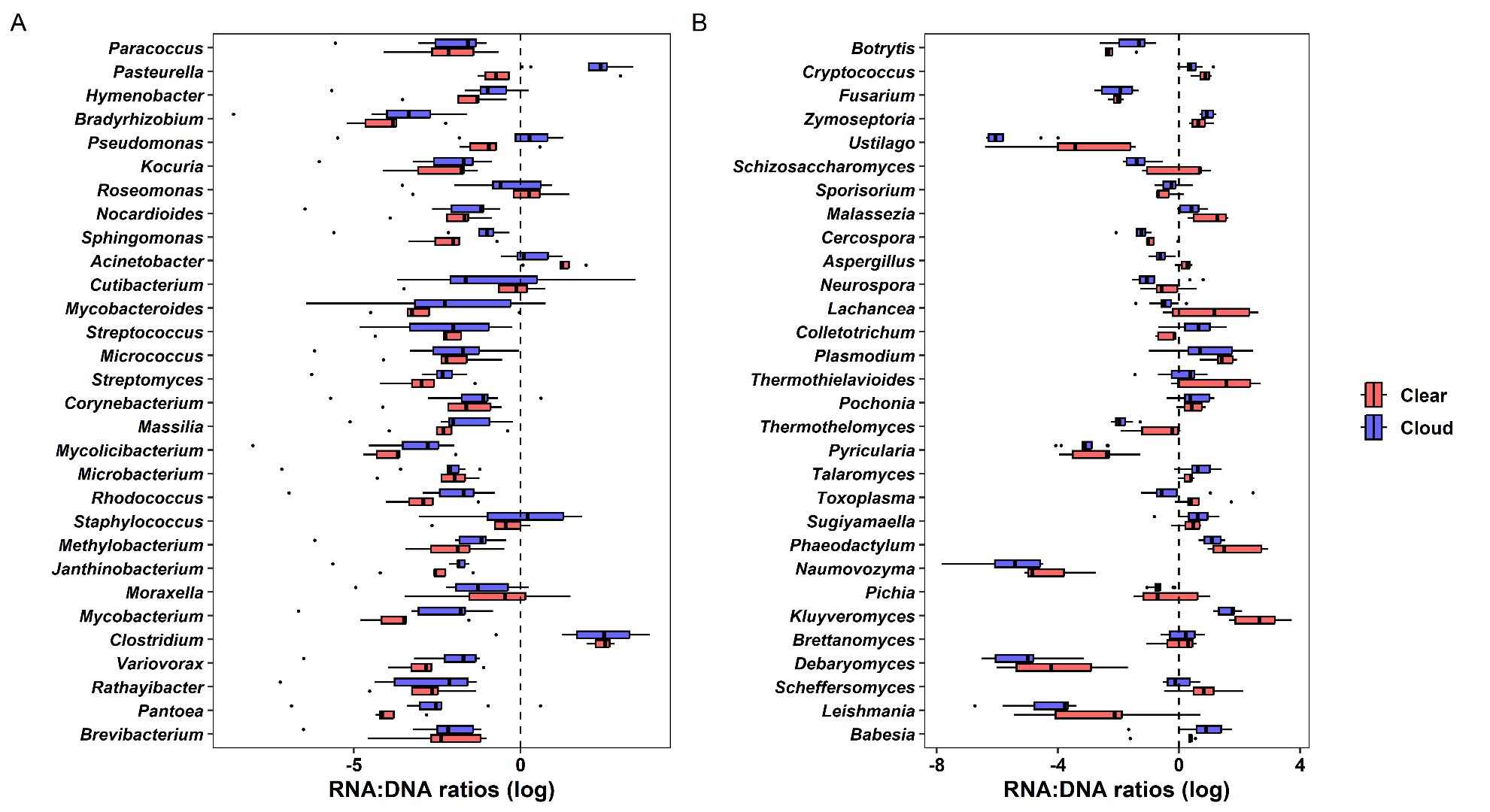
**Fig. S8.**

Relative representation of read numbers affiliated with genera of (A) the 30 most represented bacteria, and (B) eukaryotes, in MTs *versus* MGs, for cloudy and clear atmospheric conditions. Taxa are ordered from top to bottom in descending order of the number of reads in MGs.

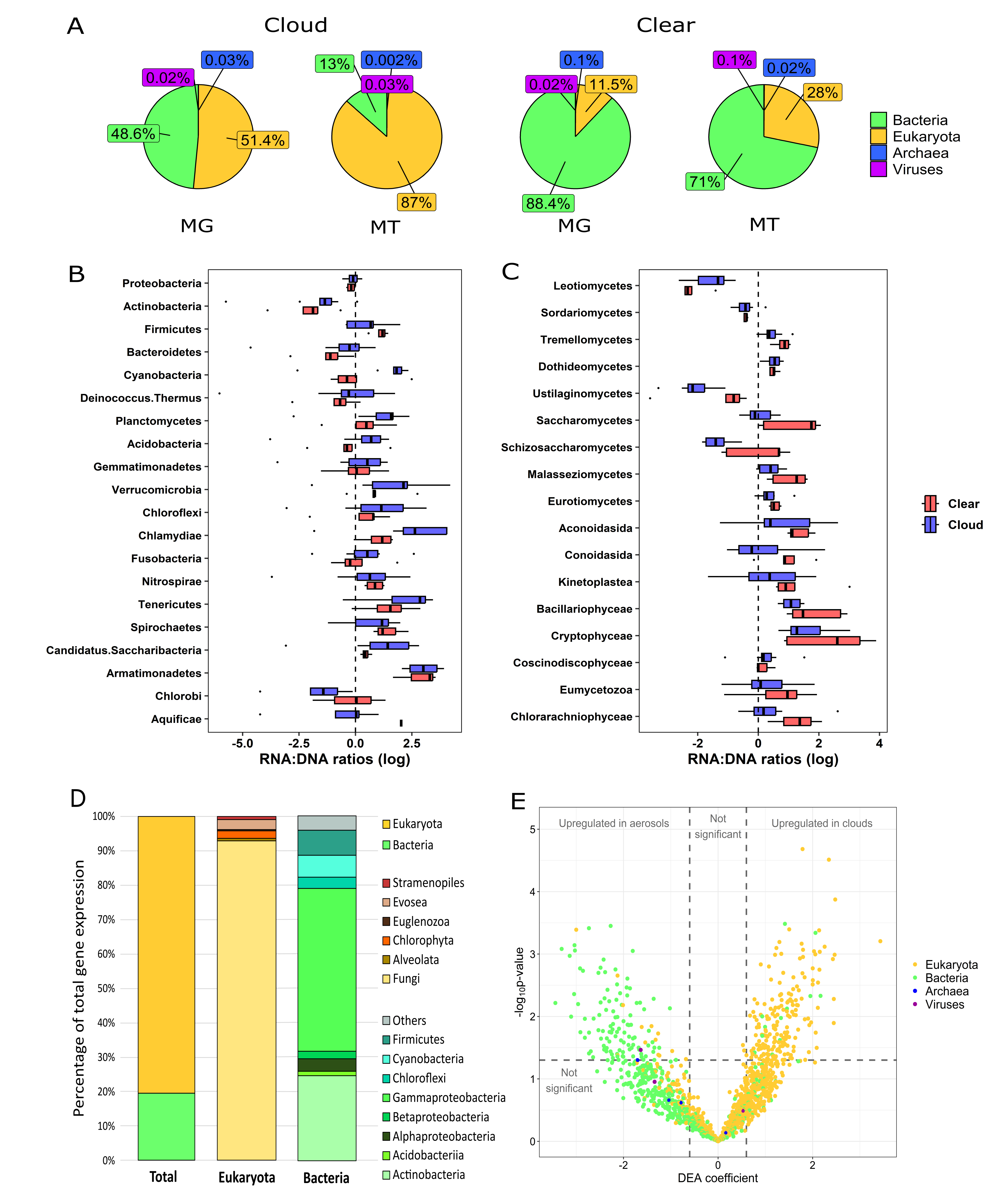

**Fig. S9.**

Proportion (%) of overexpressed genes affiliated to Eukaryotes or Bacteria and their respective phyla (or classes of Proteobacteria, *i.e.*, Pseudomonadota), based on the 488 overexpressed genes detected by differential expression analysis.

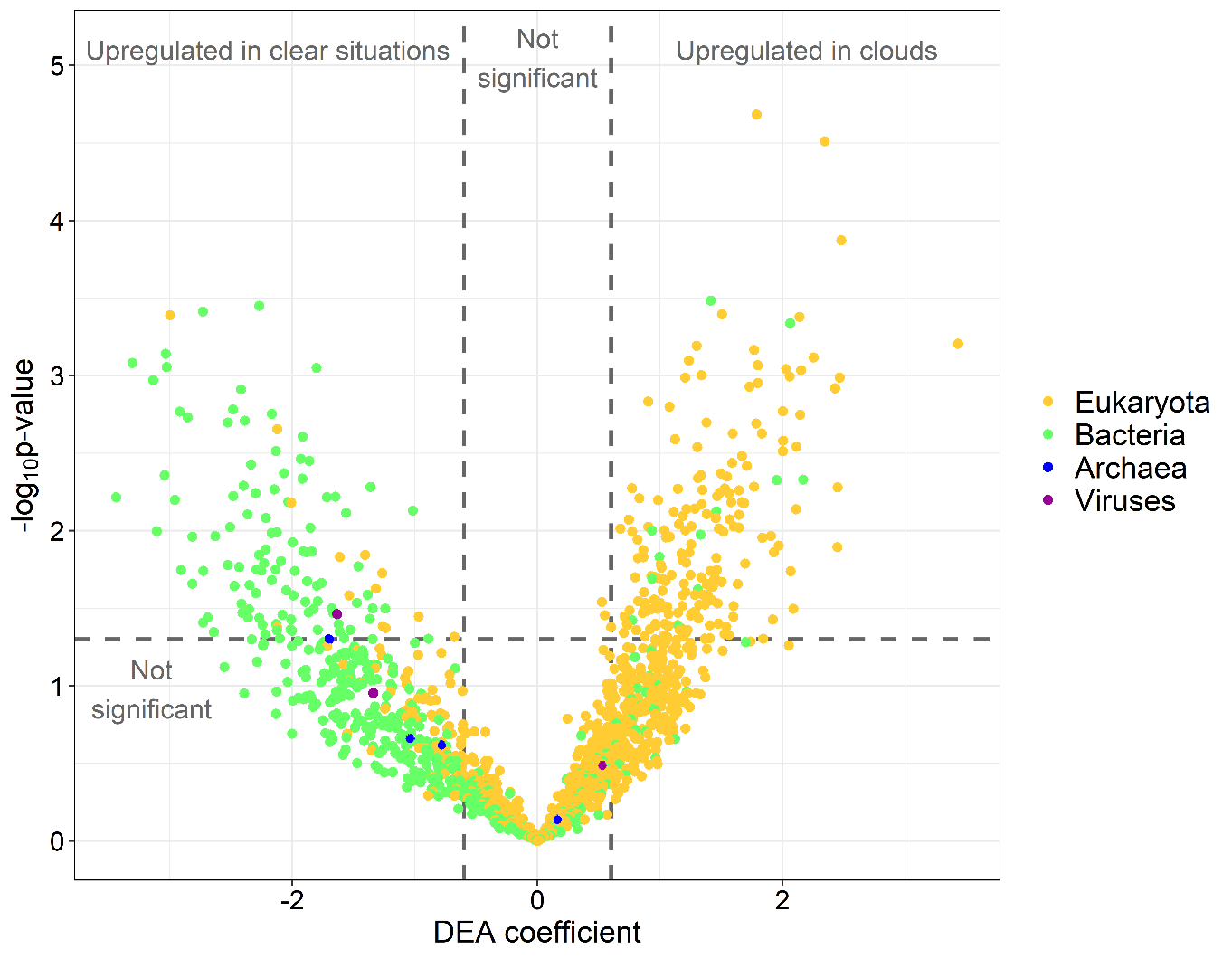

**Fig. S10.**

Volcano plot representing differential gene expression in clouds (positive coefficients) compared to clear conditions (negative coefficients), colored by kingdom; Dashed lines indicate significance thresholds.

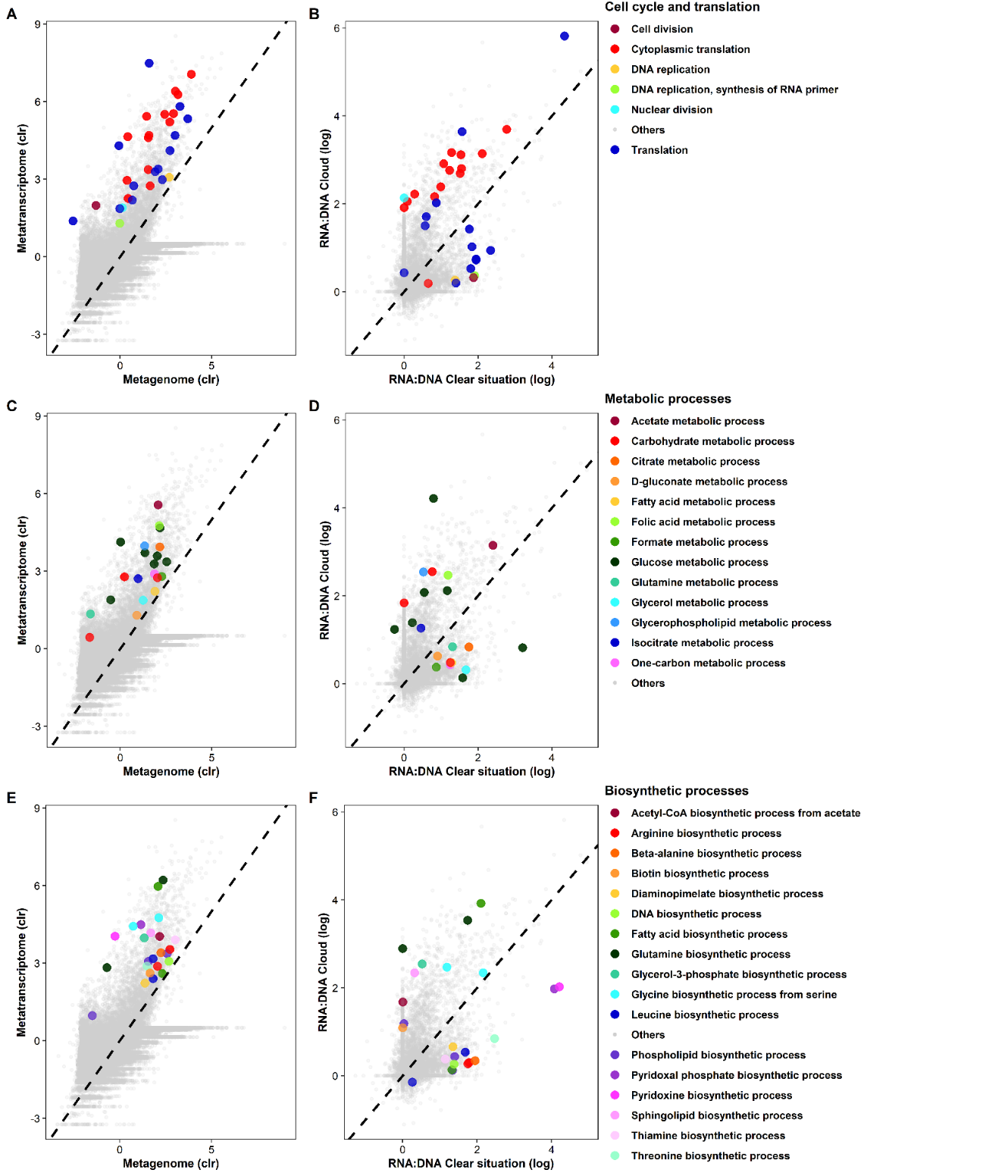

Fig. S11.

Relative representation of Gene Ontology terms (GOs) in MTs *versus* MGs, all samples considered (A; C; E), and functional expression levels in clouds *versus* clear conditions based on GO representation in MTs *versus* MGs, termed as RNA:DNA (B; D; F), for Biological Processes GOs related to cell cycle and translation (A; B), metabolic processes (C; D), and biosynthetic processes (E; F); clr: centered-log ratio transformation.

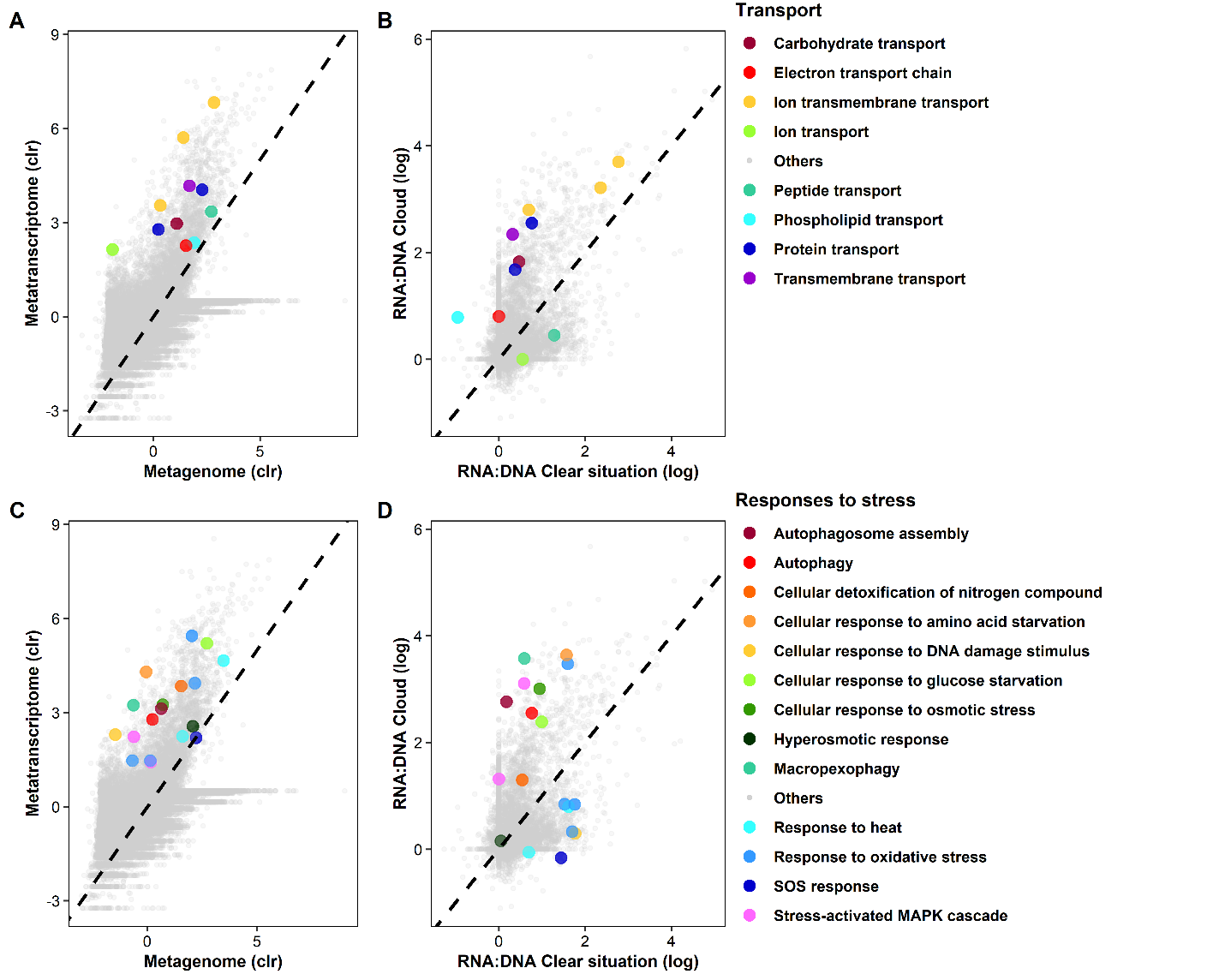
Fig. S12.

Same as Fig. S7, for Biological Processes GOs related to transport (A; B) and responses to stress (C; D).

Table S1.

Nucleic acid concentrations in the air volumes sampled during cloudy and clear conditions, and corresponding numbers of annotated genes.

| **Sample ID** | **Total DNA concentration**  **(ng.m^-3^ of air)** ^#^ | **Total RNA concentration**  **(ng.m^-3^ of air)** ^#^ | **RNA:DNA concentrations ratio** | **Number of annotated genes in MG** | **Number of annotated genes in MT** |
| --- | --- | --- | --- | --- | --- |
| **CLEAR CONDITIONS** |  |  |  |  |  |
| 20200707AIR | 2.19 | 0.47 | 0.21 | 5,463 | 2,237 |
| 20200708AIR | 1.35 | 0.38 | 0.28 | 3,287 | 2,095 |
| 20200709AIR | 0.87 | 0.33 | 0.38 | 6,649 | 1,676 |
| 20200922AIR | 0.70 | 0.68 | 0.98 | 11,690 | 2,248 |
| 20201118AIR | 0.14 | 0.23 | 1.64 | 15,431 | 4,471 |
| 20201124AIR | 0.12 | 0.06 | 0.53 | - | - |
| **Minimum** | **0.12** | **0.06** | **0.21** | **3,287** | **1,676** |
| **Maximum** | **2.19** | **0.68** | **1.64** | **15,431** | **4,471** |
| **Median** | **0.78** | **0.36** | **0.46** | **6,649** | **2,237** |
| **Mean** | **0.89** | **0.36** | **0.67** | **8,504** | **2,545** |
| **Standard error** | **0.78** | **0.21** | **0.55** | **4,951** | **1,101** |
| **CLOUDS** |  |  |  |  |  |
| 20191002CLOUD | 0.18 | 0.32 | 1.77 | 19,010 | 2,219 |
| 20191022CLOUD | 0.24 | 0.65 | 2.70 | 14,804 | 3,855 |
| 20200311CLOUD | 0.31 | 0.62 | 1.98 | 14,220 | 985 |
| 20200717CLOUD | 0.33 | 0.52 | 1.59 | 12,491 | 1,477 |
| 20201016CLOUD | 0.16 | 0.23 | 1.49 | 17,912 | 6,134 |
| 20201028CLOUD | 0.16 | 0.36 | 2.24 | 16,271 | 3,368 |
| 20201103CLOUD | 0.37 | 0.97 | 2.65 | 15,958 | 2,737 |
| 20201110CLOUD | 0.38 | 1.27 | 3.38 | 16,527 | 3,110 |
| 20201119CLOUD | 0.28 | 1.00 | 3.62 | 16,064 | 3,406 |
| **Minimum** | **0.16** | **0.23** | **1.49** | **12,491** | **985** |
| **Maximum** | **0.38** | **1.27** | **3.62** | **19,010** | **6,134** |
| **Median** | **0.28** | **0.62** | **2.24** | **16,064** | **3,110** |
| **Mean** | **0.27** | **0.66** | **2.38** | **15,917** | **3,032** |
| **Standard error** | **0.09** | **0.35** | **0.76** | **1,934** | **1,496** |
| **P-value** (Mann-Whitney test; cloudy *vs* non-cloudy conditions) | **0.32** | **0.16** | **0.004**** | **0.01*** | **0.59** |

^#^: as inferred from quantification in the extracts, based on sampling time and flow rate; NA: no data available; *: significant p-value (< 0.05); **: highly significant p-value (< 0.01).

Table S2.

Processing information of sequences in MGs for (A) cloudy and (B) clear conditions.

| **A)** | CLOUD  20191002 | CLOUD  20191022 | CLOUD  20200311 | CLOUD  20200717 | CLOUD  20201016 | CLOUD  20201028 | CLOUD  20201103 | CLOUD  20201110 | CLOUD  20201119 |
| --- | --- | --- | --- | --- | --- | --- | --- | --- | --- |
| Number of raw reads | 65,812,666 | 259,998,456 | 58,746,330 | 43,757,944 | 97,184,944 | 66,400,646 | 54,975,726 | 60,932,548 | 77,768,050 |
| After quality control (QC) | 41,675,340 | 175,692,186 | 40,113,524 | 29,200,252 | 64,669,674 | 42,495,176 | 35,964,782 | 40,917,502 | 51,587,742 |
| % removed | 37 | 32 | 32 | 33 | 33 | 36 | 35 | 33 | 34 |
| Number of rRNA gene reads | 479,510 | 2,105,871 | 681,376 | 497,119 | 825,886 | 553,734 | 461,444 | 569,711 | 608,520 |
| % of rRNA gene reads | 1.15 | 1.20 | 1.70 | 1.70 | 1.28 | 1.30 | 1.28 | 1.39 | 1.18 |
| Number of non-rRNA gene reads | 41,195,830 | 173,586,315 | 39,432,148 | 28,703,133 | 63,843,788 | 41,941,442 | 35,503,338 | 40,347,791 | 50,979,222 |
| % of non-rRNA gene reads | 98.85 | 98.80 | 98.30 | 98.30 | 98.72 | 98.70 | 98.72 | 98.61 | 98.82 |
| % of human reads in non-rRNA gene reads | 0.11 | 0.71 | 0.02 | 0.04 | 0.08 | 0.14 | 0.07 | 0.08 | 0.09 |
| Number of assembled contigs | 194,547 | 495,663 | 275,249 | 129,639 | 316,632 | 213,954 | 193,399 | 241,377 | 207,227 |
| Number of properly paired reads | 2,060,240 | 17,067,006 | 3,109,058 | 1,853,656 | 5,065,598 | 3,131,000 | 3,707,712 | 3,154,518 | 2,198,260 |
| % of properly mapped reads | 5 | 9.9 | 7.9 | 6.4 | 7.9 | 7.5 | 10.4 | 7.8 | 4.3 |
| Number of affiliated reads | 554,018 | 2,271,135 | 480,815 | 506,200 | 769,566 | 469,334 | 337,543 | 374,346 | 597,562 |
| % affiliated | 2.66 | 2.59 | 2.40 | 3.47 | 2.38 | 2.21 | 1.18 | 1.83 | 2.32 |
| Number of affiliated human reads | 41,379 | 654,422 | 19,638 | 19,244 | 38,655 | 35,478 | 18,311 | 26,211 | 39,429 |
| % of human reads | 7.5 | 28.8 | 4.1 | 3.8 | 5 | 7.6 | 5.4 | 7 | 6.6 |

| **B)** | CLEAR  20200707 | CLEAR  20200708 | CLEAR  20200709 | CLEAR  20200922 | CLEAR  20201118 | CLEAR  20201124 |
| --- | --- | --- | --- | --- | --- | --- |
| Number of raw reads | 47,409,014 | 30,435,818 | 44,295,022 | 41,909,288 | 40,730,342 | 41,086,722 |
| After quality control (QC) | 31,161,586 | 19,145,910 | 30,985,400 | 31,152,980 | 25,543,444 | 28,294,184 |
| % removed | 34 | 37 | 30 | 26 | 37 | 31 |
| Number of rRNA gene reads | 524,095 | 361,927 | 333,292 | 437,338 | 384,484 | 305,043 |
| % of rRNA gene reads | 1.68 | 1.89 | 1.08 | 1.40 | 1.51 | 1.08 |
| Number of non-rRNA gene reads | 30,637,491 | 18,783,983 | 30,652,108 | 30,715,642 | 25,158,960 | 27,989,141 |
| % of non-rRNA gene reads | 98.32 | 98.11 | 98.92 | 98.60 | 98.49 | 98.92 |
| % of human reads in non-rRNA gene reads | 0.02 | 0.02 | 0.01 | 0.02 | 0.14 | - |
| Number of assembled contigs | 140,392 | 43,627 | 99,268 | 179,815 | 101,745 | - |
| Number of properly paired reads | 4,511,490 | 2,149,274 | 3,511,940 | 5,361,278 | 1,450,314 | - |
| % of properly mapped reads | 14.6 | 11.3 | 11.4 | 17.4 | 5.8 | - |
| Number of affiliated reads | 628,977 | 361,163 | 614,940 | 303,720 | 499,417 | - |
| % affiliated | 4.04 | 3.77 | 3.97 | 1.95 | 3.91 | - |
| Number of affiliated human reads | 48,092 | 27,287 | 14,197 | 23,093 | 36,927 | - |
| % of human reads | 7.6 | 7.6 | 2.3 | 7.6 | 7.4 | - |

Table S3.

Processing information of sequences in MTs for (A) cloudy and (B) clear conditions.

| **A)** | CLOUD  20191002 | CLOUD  20191022 | CLOUD  20200311 | CLOUD  20200717 | CLOUD  20201016 | CLOUD  20201028 | CLOUD  20201103 | CLOUD  20201110 | CLOUD  20201019 |
| --- | --- | --- | --- | --- | --- | --- | --- | --- | --- |
| Number of raw reads | 93,499,990 | 186,010,124 | 82,131,152 | 81,247,584 | 79,489,694 | 85,009,094 | 110,129,198 | 69,916,188 | 195,503,428 |
| After quality control (QC) | 64,702,194 | 94,221,480 | 61,733,384 | 60,764,224 | 54,020,462 | 64,420,980 | 71,515,884 | 50,964,222 | 132,952,184 |
| % removed | 31 | 49 | 25 | 25 | 32 | 24 | 35 | 27 | 32 |
| Number of rRNA reads | 59,259,226 | 85,291,849 | 58,058,408 | 56,202,145 | 43,175,946 | 57,199,748 | 28,842,768 | 45,243,381 | 115,553,819 |
| % of rRNA reads | 91.59 | 90.52 | 94.05 | 92.49 | 79.93 | 88.79 | 40.33 | 88.77 | 86.91 |
| Number of non-rRNA reads | 5,442,968 | 8,929,631 | 3,674,976 | 4,562,079 | 10,844,516 | 7,221,232 | 42,673,116 | 5,720,841 | 17,398,365 |
| % of non-rRNA reads | 8.41 | 9.48 | 5.95 | 7.51 | 20.07 | 11.21 | 59.67 | 11.23 | 13.09 |
| % of human reads in non-rRNA reads | 0.86 | 0.05 | 0.32 | 0.42 | 0.82 | 0.46 | 89.47 | 0.29 | 0.31 |
| Number de reads properly paired | 275,906 | 655,764 | 262,850 | 275,216 | 833,402 | 697,632 | 464,440 | 423,202 | 1,007,468 |
| % of properly mapped reads | 4.6 | 7 | 7 | 5.7 | 7.5 | 8.9 | 3.9 | 7.1 | 5.6 |
| Number of affiliated reads | 24,817,282 | 37,049,964 | 26,829,087 | 22,589,666 | 18,755,564 | 24,842,334 | 32,332,229 | 17,680,329 | 52,249,268 |
| % affiliated | 76.71 | 78.64 | 86.92 | 74.35 | 69.44 | 77.12 | 90.42 | 69.38 | 78.60 |
| Number of affiliated human reads | 31,612 | 9,229 | 15,483 | 26,700 | 64,652 | 25,003 | ≈,19,000,000 | 22,925 | 46,348 |
| % of human reads | 0.1 | 0.02 | 0.06 | 0.1 | 0.3 | 0.1 | ≈,59 | 0.1 | 0.1 |

| **B)** | CLEAR  20200707 | CLEAR  20200708 | CLEAR  20200709 | CLEAR  20200922 | CLEAR  20201118 | CLEAR  20201124 |
| --- | --- | --- | --- | --- | --- | --- |
| Number of raw reads | 71,487,464 | 116,985,074 | 96,661,644 | 65,554,022 | 76,813,684 | 68,355,146 |
| After quality control (QC) | 49,784,022 | 63,268,244 | 56,706,232 | 48,733,804 | 54,728,858 | 54,728,858 |
| % removed | 30 | 46 | 41 | 26 | 29 | 20 |
| Number of rRNA reads | 41,199,078 | 51,568,599 | 49,530,808 | 42,808,640 | 45,507,747 | 5,986,089 |
| % of rRNA reads | 82.76 | 81.51 | 87.35 | 87.84 | 83.15 | 12.06 |
| Number of non-rRNA reads | 8,584,944 | 11,699,645 | 7,175,424 | 5,925,164 | 9,221,111 | 43,642,337 |
| % of non-rRNA reads | 17.24 | 18.49 | 12.65 | 12.16 | 16.85 | 87.94 |
| % of human reads in non-rRNA reads | 1.95 | 1.37 | 12.02 | 0.2 | 0.7 | - |
| Number of reads properly paired | 648,418 | 383,000 | 190,608 | 667,098 | 467,706 \| | - |
| % of properly mapped reads | 7.6 | 3.3 | 2.7 | 10.8 | 4.9 | - |
| Number of affiliated reads | 18,903,493 | 22,757,792 | 22,429,335 | 19,290,161 | 17,432,686 | - |
| % affiliated | 75.94 | 71.94 | 79.11 | 79.17 | 63.71 | - |
| Number of affiliated human reads | 103,756 | 119,150 | 164,072 | 18,204 | 96,729 | - |
| % of human reads | 0.5 | 0.5 | 0.7 | 0.1 | 0.6 | - |

**Other Supplementary Materials (separate electronic files):**

- **Data S1.** Read counts affiliated with Bacteria in MGs based on taxonomy annotations in Kraken’s plusPF database, at various taxonomic levels. The samples are named as “A” for clear conditions or “C” for clouds, respectively, followed by the sampling date as “yyyymmdd”.
- **Data S2.** Read counts affiliated with Eukaryota in MGs based on taxonomy annotations in Kraken’s plusPF database, at various taxonomic levels. The samples are named as “A” for clear conditions or “C” for clouds, respectively, followed by the sampling date as “yyyymmdd”.
- **Data S3.** Taxon-based differential expression analysis (DEA), from bacterial and Eukaryotic families and genera representation in MTs *versus* MGs, all samples considered without distinction between atmospheric conditions. Positive coefficients highlighted in green indicate taxa significantly more represented in MTs than in MGs. Value: factor of reference; coef: DEA coefficient from MTXmodel R Package; stderr: standard deviation of the DEA coefficient; N: total number of samples considered; N.not.0: number of samples with corresponding reads number >0; pval: p-value associated with the DEA coefficient.
- **Data S4.** List of the 488 overrepresented transcripts based on differential expression analysis (DEA), without distinction between atmospheric conditions, and proteins, genes, organisms, GO terms and E.C. numbers associated with them. Value: factor of reference for which positive DEA coefficients indicate overrepresentation; coef: DEA coefficient from MTXmodel; stderr: standard deviation of the DEA coefficient; N: total number of samples considered; N.not.0: number of samples with corresponding reads number >0; pval: p-value associated with the DEA coefficient.
- **Data S5.** Average DEA coefficients of GO terms associated with overrepresented transcripts, all samples considered without distinction between atmospheric conditions. Positive coefficients are highlighted in green and indicate overrepresented GO terms.
- **Data S6.** List of the 320 transcripts differentially represented between clouds and clear atmosphere, based on differential expression analysis (DEA), and proteins, genes, organisms, GO terms and E.C. numbers associated with them. Value: factor of reference for which positive DEA coefficients indicate overrepresentation; coef: DEA coefficient from MTXmodel; stderr: standard deviation of the DEA coefficient; N: total number of samples considered; N.not.0: number of samples with corresponding reads number >0; pval: p-value associated with the DEA coefficient.
- **Data S7.** Average DEA coefficients of GO terms associated with differentially represented transcripts between clouds and clear atmosphere. Positive coefficients are highlighted in green and indicate GO terms overrepresented in clouds.
